# Constitutive PI3K–AKT activation promotes steatohepatitis through *de novo* lipogenesis despite enhanced mitochondrial β-oxidation, essential fatty acid depletion and reduced lipid peroxidation

**DOI:** 10.64898/2026.09.23.753762

**Authors:** Albert S. Peixoto, Érique Castro, Bianca F. Leonardi, Tiago E. Oliveira, Adriano B. Chaves-Filho, Thayna S. Vieira, Caroline A. Tomazelli, Natalia Monteiro Pessoa, Erika Vicência Monteiro Pessoa, Marina A. Abe-Honda, Ana B. Pires, Luciano P. Silva Júnior, Erick M. Silveira, Beatriz Pereira da Silva, Camila Nogueira, Flavia Carla Meotti, Marcos Y. Yoshinaga, Sayuri Miyamoto, William T. Festuccia

## Abstract

**Background.:** Activation of *de novo* lipogenesis (DNL) is a major hallmark of steatotic liver disease (SLD), a highly prevalent group of diseases ranging from a harmless steatosis to more severe steatohepatitis, cirrhosis, and hepatocellular carcinoma. We interrogated herein the early alterations in liver proteome, transcriptome, lipidome, inflammation, fibrosis, and oxidative status in a mouse model of progressive SLD induced by constitutive activation of PI3K-AKT signaling, the major inducer of DNL.

**Methods.:** Male, 8 weeks-old mice bearing phosphatase and tensin homolog (*Pten*) deletion and therefore constitutive activation of PI3K-AKT in hepatocytes (albumin-Cre) and littermate controls were evaluated for liver steatosis, injury, lipid peroxidation, metabolism, proteome, lipidome and transcriptome either upon feeding with a chow or high-fat diets supplemented or not with linoleic acid or after treatment with vehicle or acetyl-CoA carboxylase (ACC) inhibitor (ND-630, 20 mg/kg/day, i.p.)

**Results.:** Mice bearing *Pten* deletion and therefore constitutive activation of PI3K-AKT in hepatocytes display steatohepatitis characterized by enhanced DNL, severe steatosis, hepatocyte ballooning, inflammation, fibrosis and hepatomegaly, which, unexpectedly, was associated with reduced lipid peroxidation, increased GSH content and enhanced rates of triacylglycerol turnover, fatty acid β-oxidation, tricarboxylic acid cycle flux, and mitochondrial respiration, altogether promoting a robust lipidome remodeling mainly defined by enrichment of DNL-derived fatty acids in detriment of a broad depletion of essential fatty acids (linoleic and α-linolenic acids) and cardiolipins. Dietary restoration of liver linoleic acid content mildly attenuated liver inflammation, without however affecting lipid peroxidation and improving steatosis and fibrosis, whereas pharmacological acetyl-CoA carboxylase (ACC) and DNL inhibition markedly reduced steatosis, inflammation and fibrosis, despite further lowering hepatic linoleic acid content.

**Conclusions.:** Collectively, these findings support a harmful role of essential fatty acids depletion fomenting liver inflammation and identify DNL as a major culprit of steatohepatitis induced by constitutive PI3K-AKT activation.

**What is already known on this topic:**

- *Pten* inactivation in hepatocytes leading to constitutive PI3K-AKT signaling and *de novo* lipogenesis activation (DNL) is a recurrent liver signature found in steatotic liver disease (SLD)-bearing patients, promoting disease progression through not completely defined molecular mechanisms.

**What this study adds:**

- *Pten* inactivation in hepatocytes promotes liver steatosis, injury, inflammation and fibrosis, in spite of depleting essential fatty acids and cardiolipin, reducing lipid peroxidation, enhancing antioxidant capacity and mitochondrial mass and respiration, and eliciting lipid-related futile metabolic cycles.
- Dietary restoration of liver essential linoleic acid partially attenuates inflammation, but not steatosis and fibrosis induced by *Pten* inactivation in hepatocytes
- Pharmacological ACC and DNL inhibition mitigate the steatosis, inflammation and fibrosis induced by *Pten* inactivation in hepatocytes, but further deplete liver essential fatty acids.

**How this study might affect research, practice or policy:**

- This study raises evidence supporting a harmful role of essential fatty acid depletion exacerbating liver inflammation.
- Pharmacological strategies that increase fatty acid oxidation including ACC inhibition may enhance, while dietary supplementation may mitigate essential fatty acid depletion.
- Despite protecting healthy cells from oxidative damage, the enhanced antioxidant capacity induced by *Pten* inactivation in hepatocytes may favor survival of hepatocytes bearing DNA mutations and tumorigenesis.

## 1. Introduction

Steatotic liver disease (SLD) is a highly prevalent spectrum of hepatic diseases encompassing from a harmless steatosis to more severe steatohepatitis, cirrhosis, and hepatocellular carcinoma (HCC) [1,2]. These diseases share hepatocyte lipid accumulation and remodeling as main features, driven in part by enhanced *de novo* lipogenesis (DNL) [3–6]. DNL refers to the conversion of cytosolic citrate into palmitic acid (PA, 16:0) sequentially catalyzed by ATP citrate lyase (ACLY), acetyl-CoA carboxylase (ACC), and fatty acid synthase (FAS) [7], a metabolic pathway mainly stimulated by insulin via PI3K-AKT signaling [8]. Palmitic acid (16:0) can then be either desaturated and/or elongated into palmitoleic (POA; 16:1n7), stearic (SA; 18:0), and oleic (OA; 18:1n9) acids through reactions involving stearoyl-CoA desaturase 1 (SCD1) and fatty acid elongases [9]. Because mammals cannot synthesize the ω-6 (n-6) and ω-3 (n-3) polyunsaturated fatty acids (PUFAs) linoleic (LA; 18:2n-6) and α-linolenic (ALA; 18:3n-3) acids, respectively, these fatty acids are essential and must be acquired from diet. LA can then be elongated and desaturated to arachidonic acid (AA; 20:4n-6), while ALA can be elongated and desaturated to eicosapentaenoic (EPA; 20:5n-3) and docosahexaenoic (DHA; 22:6n-3) acids, all of which serving as precursors for the synthesis of important lipid mediators [10,11]. Notably, AA esterified in phosphatidylethanolamine is a major substrate for enzymatic, iron-dependent lipid peroxidation forming phospholipid hydroperoxides, which in excess triggers ferroptosis, an iron-dependent form of regulated cell death [12].

Liver lipid remodeling promoted by enhanced DNL is characterized by enrichment of DNL-synthesized fatty acids [13,14], which unexpectedly occurs along with a robust depletion of the essential PUFAs LA and ALA and their elongated products AA, EPA and DHA [5,15–17]. The underlying mechanisms and impact of such essential PUFA depletion on SLD progression are unknown, with studies showing conflicting findings. Indeed, higher liver LA and AA content was associated with reduced inflammation and risk of hepatic steatosis and fibrosis [18,19], while dietary LA supplementation reduced liver lipid content without increasing inflammation [20]. Conversely, intake of a LA-enriched diet increased membrane LA content, lipid peroxidation, and mitochondrial DNA damage in mice [21].

In addition to fatty acids synthesis, DNL also regulates hepatic lipid metabolism [22,23] via post-translational protein acetylation, malonylation, and palmitoylation, while the DNL intermediate malonyl-CoA allosterically inhibits carnitine palmitoyltransferase 1α (CPT1α), limiting mitochondrial fatty acid entry and β-oxidation [22]. Paradoxically, despite CPT1α inhibition, DNL activation is consistently associated with an upregulation of fatty acid β-oxidation in SLD patients [24–27], including in those with HCC [28], but also in mice bearing SLD due to severe lipoatrophy [29]. Interestingly, pharmacological peroxisome proliferator-activated receptor α (PPARα) activation enhances liver fatty acid synthesis and oxidation, and reduces hepatic LA content and inflammation in mice [30], suggesting a PPARα role in mediating these SLD phenotypes. To further interrogate the impact of constitutive DNL activation on SLD progression, we characterized herein the early alterations in liver metabolism, lipidome, proteome, transcriptome, inflammation, fibrosis, and redox status in 8 weeks old male mice featuring or not *Pten* deletion in hepatocytes exposed to either fasting/refeeding, or dietary LA supplementation, or pharmacological ACC inhibition. Noteworthy, hepatocyte PTEN inactivation, a recurrent signature found in SLD patients [31,32], promotes a temporally defined SLD progression from steatosis to steatohepatitis and HCC, serving therefore as a robust model to investigate the impact of constitutive PI3K-AKT and DNL activation on liver disease and associated lipid remodeling.

## 2. Materials and Methods

### 2.1. Mice

Mice experimental procedures were approved by the Animal Care Committee of the Institute of Biomedical Sciences, University of Sao Paulo (#115/2016 and 6160250820, CEUA). Only male mice on a C57BL/6J background were used. Mice with Pten deletion in hepatocytes (*Pten*^Lox/Lox^; *albumin*-*cre*^+/-^, referred henceforth as L-PtenKO mice) and littermate controls (*Pten*^Lox/Lox^, referred henceforth as L-PtenWT) were produced by crossing *Pten*^Lox/Lox^ mice (B6.129S4-*Pten^tm1Hwu^*/J, Jackson Laboratories) with *albumin*-*cre* mice (B6.Cg-*Speer6-ps1^Tg(Alb-cre)21Mgn^*/J, Jackson Laboratories) as described previously [33]. Mouse genotype was determined by PCR analysis of tail genomic DNA, and Pten deletion was confirmed by Western blotting. 8 weeks old, male mice kept at 23 ± 1°C on a 12:12 h light-dark cycle (lights on at 06:00 h), fed a nonpurified chow diet (70% carbohydrate, 20% protein, 10% fat, in % kcal, NUVILAB CR-1®-Sogorb Inc., Paraná, Brazil) were euthanized after either 10 h of fasting (from 22:00 to 08:00 h) or 10 h of fasting followed by 3 h of refeeding by exsanguination after anesthesia with isoflurane. A separate cohort of chow-fed L-PtenWT and L-PtenKO mice was treated with either vehicle (1% DMSO in 0.2% carboxymethylcellulose) or the acetyl-CoA carboxylase (ACC) inhibitor ND-630 (20 mg/kg/day, i.p.; InvivoChem) [34] for 14 days and euthanized after 10 h of fasting (22:00-08:00 h) followed by 3 h of refeeding. Another cohort of 8 weeks old L-PtenWT and L-PtenKO mice was fed either a purified high-carbohydrate diet (70% carbohydrate, 20% protein, and 10% fat) containing 14.4 g/kg of linoleic acid [18:2n6] (LF) or high-fat diets (40% carbohydrate, 20% protein, and 40% fat) containing either 41 g/kg of linoleic acid (HF) or 77.3 g/kg of linoleic acid (HFLA) for 4 weeks. Mice were euthanized after 10 h of fasting (22:00-08:00 h), followed by 3 h of refeeding. Diet compositions are detailed in Supplementary Table 1

### 2.2. Glucose and lipid metabolism in liver slices

Liver slices (1000 μm thick) produced with a McIlwain Tissue Chopper were incubated in hermetically closed vials containing 1.5 mL of Krebs-Ringer bicarbonate buffer (pH 7.4) composed of (in mM) 5 glucose, 0.51 MgCl₂, 4.56 KCl, 119.8 NaCl, 0.7 Na₂HPO₄, 1.3 NaH₂PO₄, and 15.0 NaHCO₃, supplemented with 1% essentially fatty acid-free albumin and containing either 0.3 μCi/vial of [U-¹⁴C]glucose for evaluation of glucose metabolism (oxidation and conversion to lactate, glycerol, and fatty acids from triacylglycerol and glycogen), 2 mM acetate and 0.3 μCi/vial of [1-¹⁴C]acetate for evaluation of fatty acid synthesis (*de novo* lipogenesis) and oxidation (tricarboxylic acid cycle flux), 100 μM palmitic acid and 0.3 μCi/vial of [1-¹⁴C]palmitic acid for evaluation of palmitic acid oxidation and incorporation into triacylglycerol, or 100 μM linoleic acid and 0.3 μCi/vial of [1-¹⁴C]linoleic acid for evaluation of linoleic acid oxidation and incorporation into triacylglycerol. After 2 h at 37°C, 50 μL of medium was collected for lactate measurement (labeled glucose only), and the medium was acidified for collection of labeled CO₂ in a filter paper moistened with phenylethylamine:ethanol (1:1). Liver slices were washed with saline and destined for either glycogen extraction or lipid extraction with chloroform/methanol for measurement of acetate, palmitic acid, or linoleic acid incorporation into triacylglycerol fatty acids and glucose incorporation into triacylglycerol glycerol and fatty acids, as described [29,35]. Values were expressed per mg of protein to correct for cell number.

### 2.3. Western blotting

Liver samples were homogenized in lysis buffer containing (in mM) 50 HEPES, 40 NaCl, 50 NaF, 2 EDTA, 10 sodium pyrophosphate, 10 sodium glycerophosphate, 2 sodium orthovanadate, 1% Triton X-100, and EDTA-free protease inhibitor (Roche), centrifuged at 15,000 × g for 15 min at 4°C, and protein concentration was determined by the Bradford method [36]. Equal amounts of protein (30 μg) were separated by SDS-PAGE, transferred to PVDF membranes, blocked in TBS-T containing 5% non-fat milk for 1 h, and incubated overnight at 4°C with primary antibodies (1:1000) in TBS-T containing 5% BSA. After washing, membranes were incubated with HRP-conjugated secondary antibodies (1:5000) for 1 h, and signals were detected using enhanced chemiluminescence (ECL) on an imaging system (G:BOX, Syngene). Primary antibodies are listed in Supplementary Table 2.

### 2.4. Liver glycogen and triacylglycerol contents

Approximately 100 mg of liver was digested in 30% KOH saturated with Na₂SO₄ at 70°C. Glycogen was precipitated by centrifugation after addition of 95% ethanol. Pellets were washed, resuspended in water, hydrolyzed with 4 N H₂SO₄ at 95°C, neutralized with 4 N NaOH, and glucose levels were quantified using a colorimetric kit (Labtest, Lagoa Santa, Brazil) [37]. Hepatic triacylglycerol content was quantified in liver samples (50–70 mg) after homogenization in lysis buffer (140 mM NaCl, 50 mM Tris, pH 7.4, and 0.1% Triton X-100). Liver homogenates (10 mg/mL) were then incubated at 37°C with 1% deoxycholate for 10 min, and triacylglycerol content was measured spectrophotometrically using a colorimetric kit (Labtest, Lagoa Santa, Brazil) and normalized to wet tissue weight [38].

### 2.5. Liver cytokine and thiobarbituric acid-reactive substances (TBARS) quantification

Cytokine levels were measured in liver homogenates prepared as described in Western blotting. Approximately 50 μg of protein were used to quantify IL-6, TNF-α, IL-1β, and IL-10 using ELISA kits (DuoSET, R&D Systems, Minneapolis, MN, USA) according to the manufacturer’s instructions. Lipid peroxidation was estimated by measuring thiobarbituric acid-reactive substances (TBARS) [39]. Liver tissue was homogenized at 200 mg/mL in 50 mM Tris-HCl buffer (pH 8.0) and centrifuged at 10,000 × *g* for 20 min to obtain the supernatant. For the assay, 100 μL of liver extract or TEP standard was mixed with 100 μL of trichloroacetic acid (TCA, 10%) and 800 μL of thiobarbituric acid (TBA, 0.53%, dissolved in 20% acetic acid). Butylated hydroxytoluene (BHT, 0.02%) was added to the reaction to minimize MDA formation from sample oxidation during heating. Samples were incubated for 1 h at 95°C and then cooled on ice for 10 min. Samples were subsequently centrifuged at 5,000 × *g* for 20 min, and absorbance was measured at 535 nm. TBARS content was quantified using a standard curve generated with 1,1,3,3-tetraethoxypropane (TEP; 0–40 nmol) and expressed as MDA equivalents normalized to tissue mass (nmol MDA equivalents/mg tissue).

### 2.6. Glutathione peroxidase (GPx) activity

Approximately 100 mg of liver tissue was pulverized in liquid nitrogen and homogenized in 600 μL of 50 mM potassium phosphate buffer (pH 7.8) containing EDTA-free protease inhibitor (Roche Life Science, Pleasanton, CA, USA). Samples were sonicated (40%, 10 × 1-s pulses) and centrifuged twice at 13,000 × g for 20 min at 4°C. The supernatants were collected and used as total protein extracts for GPx activity. GPx activity was determined using an indirect coupled assay with H_2_O_2_ and cumene hydroperoxide as substrates, as previously described [40]. Briefly, 30 μL of protein extract was added to 225 μL of reaction buffer containing 55.6 mM K_2_HPO_2_·2 H_2_O (pH 7.0), 1.1 mM EDTA, 1.1 mM NaN_3_, 1.33 mM glutathione reductase, and 1.33 mM GSH, followed by 15 μL of 4 mM NADPH. Initiate the reaction by adding 30 μL of 30 mM H_2_O_2_ and monitor for 5 min at 340 nm. GPx activity was expressed as mU/mg protein.

### 2.7. Dot blotting for detection of 4-hydroxynonenal protein adducts

Liver homogenates were prepared as described above and diluted in buffer containing 50 mM HEPES, 40 mM NaCl, 50 mM NaF, 2 mM EDTA, 10 mM sodium pyrophosphate, 10 mM sodium glycerophosphate, 2 mM sodium orthovanadate, and 1% Triton X-100. Aliquots containing 2 μg of protein in 2 μL were applied to PVDF membranes. After drying, membranes were activated with methanol, blocked with 5% BSA for 1 h, and incubated overnight with anti-4-HNE (1:1,000) antibodies. Membranes were then incubated for 1 h with HRP-conjugated anti-rabbit IgG (1:5,000). Signals were detected using ECL and quantified with ImageJ. Protein loading was normalized by Coomassie Blue staining of the membranes.

### 2.8. Protein carbonylation

Protein carbonylation was assessed using the OxyBlot® Protein Oxidation Detection Kit (Merck), according to the manufacturer’s instructions.

### 2.9. Liver histology

Liver samples were fixed in 4% paraformaldehyde and embedded in paraffin. Sections 5.0 µm thick were obtained and stained with hematoxylin and eosin (H/E) to assess general liver morphology.

### 2.10. RNA extraction and qPCR

Total RNA was extracted from liver (30 mg), reverse transcribed, and used for quantitative PCR analysis as previously described [37]. Primer nucleotide sequences are detailed in Supplementary Table 3. Real-time PCR data were analyzed using the 2^-ΔΔCT method. Data are expressed as the ratio of target-gene expression to the housekeeping gene cyclophilin, whose expression was not significantly affected by mouse genotype.

### 2.11. Citrate synthase (CS), acetyl-CoA carboxylase (ACLy), fatty acid synthase (FAS) and glycerol 3-phosphate acyltransferase (GPAT) activities

For CS activity, approximately 50 mg of liver tissue was homogenized in 200 μL of buffer containing 50 mM potassium phosphate (pH 7.5) and EDTA-free protease inhibitor (Roche), and centrifuged at 13,000 × *g* for 20 min at 4°C. Enzymatic activity was measured using 10 μg of protein from the supernatant in a reaction mixture containing 60 μL of distilled water, 100 μL of 0.1 M Tris-HCl, 20 μL of 1 mM 5,5′-dithiobis-(2-nitrobenzoic acid) (DTNB), 6 μL of 10 mM acetyl-CoA, and 10 μL of 10 mM oxaloacetate. The reduction of DTNB was monitored for 5 min at 412 nm [41]. For ACLy activity, liver was homogenized in ice-cold buffer composed of 0.25 M sucrose, 2 mM EGTA, 20 mM Tris-HCl, 2% albumin, pH 7.4, and centrifuged at 400 × *g* for 5 min. The supernatant was transferred and centrifuged at 13,000 × *g* for 20 min at 4°C for isolation of the cytosolic fraction, which was evaluated for ACLy activity by measuring the disappearance of NADH at 340 nm at 25°C in a reaction buffer composed of 50 mM triethanolamine-HCl, 7 mM ATP, 10 mM potassium citrate, 0.1 mM NADH, 10 mM mercaptoethanol, 0.24 mM CoA, and NAD-malate dehydrogenase (1 unit/mL) [42]. Liver FAS activity was evaluated by monitoring NADPH oxidation at 340 nm. The assay was performed in 100 mM potassium phosphate buffer (pH 6.5) containing 0.1 mM NADPH and 25 μM acetyl-CoA, and the reaction was initiated by adding 60 μM malonyl-CoA, as described [43]. Liver total and N-ethylmaleimide (NEM)-resistant and -sensitive GPAT activities were evaluated as previously described [44].

### 2.12. Liver mitochondria isolation and respiration

Fresh livers were rapidly excised, minced, washed in ice-cold isolation buffer containing 250 mM sucrose, 10 mM HEPES, and 1 mM EGTA (pH 7.2, K⁺), and homogenized using a Potter-type homogenizer. Mitochondria were isolated by differential centrifugation: samples were centrifuged at 800 × *g* for 10 min, then at 12,000 × *g* for 10 min at 4°C. The pellet was resuspended in the same isolation buffer and centrifuged again at 12,000 × *g* for 10 min at 4°C, and the resulting pellet was resuspended in buffer containing 75 mM D-mannitol, 25 mM sucrose, 5 mM KH₂PO₄, 20 mM Tris-HCl, 100 mM KCl, 0.1% BSA, 1 mM EGTA, and 0.5 mM EDTA (pH 7.2, K⁺) [45]. Protein concentration was determined by the Bradford method [36]. Oxygen consumption of isolated mitochondria (125 μg/mL) was measured in respiration buffer containing 120 mM sucrose, 65 mM KCl, 10 mM HEPES, 2 mM MgCl₂, 1 mM KH₂PO₄, 1 mM EGTA, and 0.1% BSA (pH 7.4, K⁺) using a high-resolution respirometer (Oroboros O2k, Innsbruck, Austria), in the presence of succinate (5 mM) plus rotenone (1 μM), palmitoylcarnitine (25 μM) plus malate (2 mM), or linoleylcarnitine (25 μM) plus malate (2 mM). State 3 respiration was induced by the addition of ADP (1 mM), state 4 respiration by oligomycin (1 μg/mL), and maximal respiration by titration with 1–2 μM CCCP [29].

### 2.13. Liver fatty acid profile by gas chromatography (GC)

Fatty acid methyl ester (FAME) analysis was performed as described [46]. Briefly, 100 μL of liver homogenate (50 mg/mL in PBS (pH 7.4) containing 20 µM deferoxamine mesylate) was mixed with 1.8 mL methanol, 100 μL of acetyl chloride and 100 μL of the internal standard C13:0 (1 mg/mL of methanol), heated at 100 °C for 60 min, mixed with hexane, vortexed and centrifuged at 1500 x g for 2 min at 4 °C. The upper hexane phase was collected and evaporated, and the lipids were dissolved in 100 μL of hexane. Individual FAMEs were analyzed by gas chromatography with flame ionization detection using a Trace 1310 gas chromatograph (Thermo Scientific) equipped with a DB-FFAP column (15 m x 0.1 mm ID x 0.1 μm film thickness; J&W Scientific, Agilent Technologies). FAMEs were identified by direct comparison with a FAME standard mix (Supelco 37 Component FAME Mix; Sigma-Aldrich). Each peak was integrated, normalized to the internal standard, and expressed as nmol/g of liver.

### 2.14. Untargeted lipidomics and data processing

Lipid extraction was performed as previously described [47,48], Liver (∼ 400 mg) was homogenized in 1 mL of 10 mM phosphate buffer (pH 7.4) containing 100 μM deferoxamine mesylate. An aliquot of 250 μL of the homogenate was mixed with 550 μL methyl tert-butyl ether (MTBE) and 600 μL methanol (MeOH). Subsequently, 100 μL of this mixture was transferred and mixed with 100 μL of internal standards (10–20 ng/μL; positive and negative mixes [29]), 100 μL ultrapure water, and 300 μL ice-cold MeOH containing 10 μM butylated hydroxytoluene (BHT). After vortex mixing, add 1 mL MTBE, then incubate samples for 1 h at 20°C with agitation. Phase separation was induced by addition of 300 μL water, followed by vortexing for 30 s and incubation on ice for 10 min. Samples were centrifuged at 10,000 × g for 10 min at 4°C, and the upper organic phase (∼700 μL), containing total lipid extract, was collected, transferred to vials, and dried under nitrogen. Samples were resuspended in 100 μL isopropanol and analyzed by UHPLC (Nexera, Shimadzu, Kyoto, Japan) coupled to tandem electrospray ionization quadrupole time-of-flight mass spectrometry (ESI-Q-TOFMS; Triple TOF^®^ 6600, SCIEX, Concord, Canada), as previously described [29,48]. Lipid molecular species were identified based on retention time, accurate mass, and characteristic MS/MS fragmentation patterns, including specific fragments and/or neutral losses, using PeakView® (SCIEX, Concord, Canada) as previously described [49]. Quantification was performed by integrating the peak area of each precursor ion and normalizing it to the corresponding internal standard using MultiQuant® (SCIEX, Concord, Canada). For subsequent analysis, the relative abundance of DNL-derived fatty acids was calculated as the sum of 16:0, 16:1, 18:0, and 18:1, whereas essential fatty acids (EFAs) were calculated as the sum of 18:2n6 and 18:3n3, and PUFAs as the sum of fatty acids with chain lengths and degrees of unsaturation greater than 18:3. Lipid concentrations were expressed as nmol of lipid/mg of Tissue.

### 2.15. Targeted analysis of oxidized lipids and data processing

Oxidized lipids were extracted and analyzed by UHPLC–MS/MS exactly as previously described [29].

### 2.16. Proteomic analysis by LC–MS/MS and data processing

Liver samples were prepared for proteomic analysis using a protocol adapted from [50]. Briefly, 50 mg of liver was homogenized in 500 μL of buffer (4% SDS, 0.1 M Tris-HCl, pH 7.6, and 0.1 M DTT), centrifuged at 12,000 × g for 20 min at 4°C, and 30 μg of protein from the supernatant was separated by SDS-PAGE in 10% acrylamide gels. Proteins were then reduced with 10 mM DTT (56°C for 60 min), alkylated with 55 mM iodoacetamide (25°C for 45 min in the dark), and digested overnight with a 12.5 ng/μL trypsin solution in 50 mM ammonium bicarbonate at 37°C. Peptides were extracted twice with 30% acetonitrile/3% trifluoroacetic acid and twice with pure acetonitrile, dried in a speed-vacuum, and desalted with C18 StageTips. Analyses were performed on an Ultimate 3000 RSLCnano system coupled to an Orbitrap Fusion Lumos mass spectrometer (nanoESI-MS/MS - Thermo Scientific, RPT02H, Chagas Institute – Fiocruz) using a C18 reversed-phase analytical column with a linear gradient from 5% to 40% solvent B (95% acetonitrile in 0.1% formic acid) over 120 min at a flow rate of 250 nL/min. Full MS scans were acquired at a resolution of 120,000 over a mass range of 300–1,500 m/z. Data were acquired in data-dependent acquisition (DDA) mode, and precursor ions above an intensity threshold were selected for fragmentation by higher-energy collisional dissociation (HCD). MS/MS spectra were acquired at a resolution of 15,000, and dynamic exclusion was set to 60 s. Raw MS data were processed using MaxQuant (v2.7.3.0; Max Planck Institute of Biochemistry, Germany) [51] with label-free quantification enabled (MaxLFQ). Data were analyzed in Perseus (v2.0.10.0; Max Planck Institute of Biochemistry, Germany).[54] After removal of reverse hits, contaminants, and proteins identified only by site. LFQ intensities were log₂-transformed, median-centered, and filtered for ≥90% valid values across samples [53]. Missing values were imputed from a normal distribution (width = 0.3, downshift = 1.8). Differential expression was assessed using two-sample t-tests with permutation-based FDR < 0.05 (S₀ = 0.5). Proteins with |log₂ fold change| ≥ 0.58 were considered significantly regulated. Pathway enrichment analysis of significantly regulated proteins was performed using the WikiPathways database for mouse [54].

### 2.17. RNA sequencing and differential gene expression analysis

Total RNA was isolated from the livers of L-PtenWT and L-PtenKO mice using the RNeasy Kit (Qiagen), according to the manufacturer’s instructions. RNA samples were submitted to Azenta Life Sciences for library preparation and RNA sequencing. Poly(A)-selected RNA-seq libraries were generated and sequenced. Raw sequencing reads were processed using the Galaxy platform (usegalaxy.org) [55]. Adapter sequences and low-quality bases were removed using Trimmomatic [56], and read quality was assessed using Falco. Trimmed paired-end reads were aligned to the mouse reference genome (mm10) using HISAT2, and gene-level read counts were generated using featureCounts. The resulting count matrix was analyzed using iDEP (bioinformatics.sdstate.edu/idep/) [57]. Genes not detected across all experimental groups were excluded, and count data were normalized using the variance-stabilizing transformation (VST). Differential gene expression between L-PtenKO and L-PtenWT mice within each nutritional state was analyzed using DESeq2 [58]. Genes not detected across all experimental groups were excluded, and count data were normalized using the variance-stabilizing transformation (VST). Differential gene expression between L-PtenKO and L-PtenWT mice within each nutritional state was analyzed using DESeq2 [57]. Genes with a false discovery rate (FDR) < 0.05 and an absolute fold change > 1.5 were considered differentially expressed. We **performed pathway enrichment analysis using Enrichr** [59]**. Volcano plots were generated using GraphPad Prism** 10.4.0 (GraphPad Software, Boston, MA, USA).

### 2.18. GSH and GSSG analysis by LC–MS/MS and data processing

GSH and GSSG were extracted from 10 mg of snap-frozen liver tissue using a modified MTBE/methanol/water extraction procedure, as previously described [60]. Briefly, tissue samples were homogenized in pre-chilled methanol, followed by extraction with MTBE and water. After centrifugation, the aqueous phase was collected, dried under vacuum, and reconstituted in 50:50 (v/v) water:acetonitrile. Chromatographic separation and mass spectrometric analysis were performed by UHPLC-HILIC coupled to a Q Exactive HF-X Hybrid Quadrupole-Orbitrap mass spectrometer according to previously described conditions [61]. Briefly, metabolites were separated using an iHILIC-(P) Classic column and analyzed in negative electrospray ionization mode over an m/z range of 70–1000 at a resolution of 60,000. Peak integration was performed in Skyline (version 26), and analytes were identified by retention time and accurate mass matching within 5 ppm against an in-house library of reference standards. Peak integrations were manually inspected and corrected when necessary. Signal intensities were normalized using probabilistic quotient normalization (PQN) in MetaboAnalyst 6.0, and normalized abundances were used for subsequent statistical analyses.

### 2.19. Statistical analysis

Data are presented as mean ± SEM. Differences between two groups (L-PtenWT vs. L-PtenKO) were analyzed using an unpaired Student’s *t*-test. Two-way ANOVA followed by Tukey’s multiple comparisons test was used to assess the effects of (i) hepatocyte *Pten* deletion, fasting and refeeding, and their interaction; (ii) hepatocyte *Pten* deletion, pharmacological ACC inhibition, and their interaction; or (iii) hepatocyte *Pten* deletion, dietary linoleic acid content, and their interaction. We performed statistical analyses using GraphPad Prism (GraphPad Software, San Diego, CA, USA), and set significance at *p* ≤ 0.05.

## 3. Results

### Constitutive PI3K-AKT activation promotes steatosis, hepatomegaly, inflammation and fibrosis, despite reducing lipid peroxidation and enhancing antioxidant capacity

We successfully generated mice with hepatocyte-specific *Pten* deletion, as confirmed by the almost complete absence of PTEN in L-PtenKO livers (Fig. 1A). Loss of PTEN resulted in constitutive activation of PI3K–AKT signaling, as indicated by increased phosphorylation of AKT at Ser473 and Thr308 and of its product GSK3β at Ser9 (Fig. 1A). Despite similar body weights, L-PtenKO mice developed hepatomegaly featuring severe steatosis and increased liver mass and triacylglycerol, cholesterol, and glycogen contents (Fig. 1B–1H).

**Figure 1.**
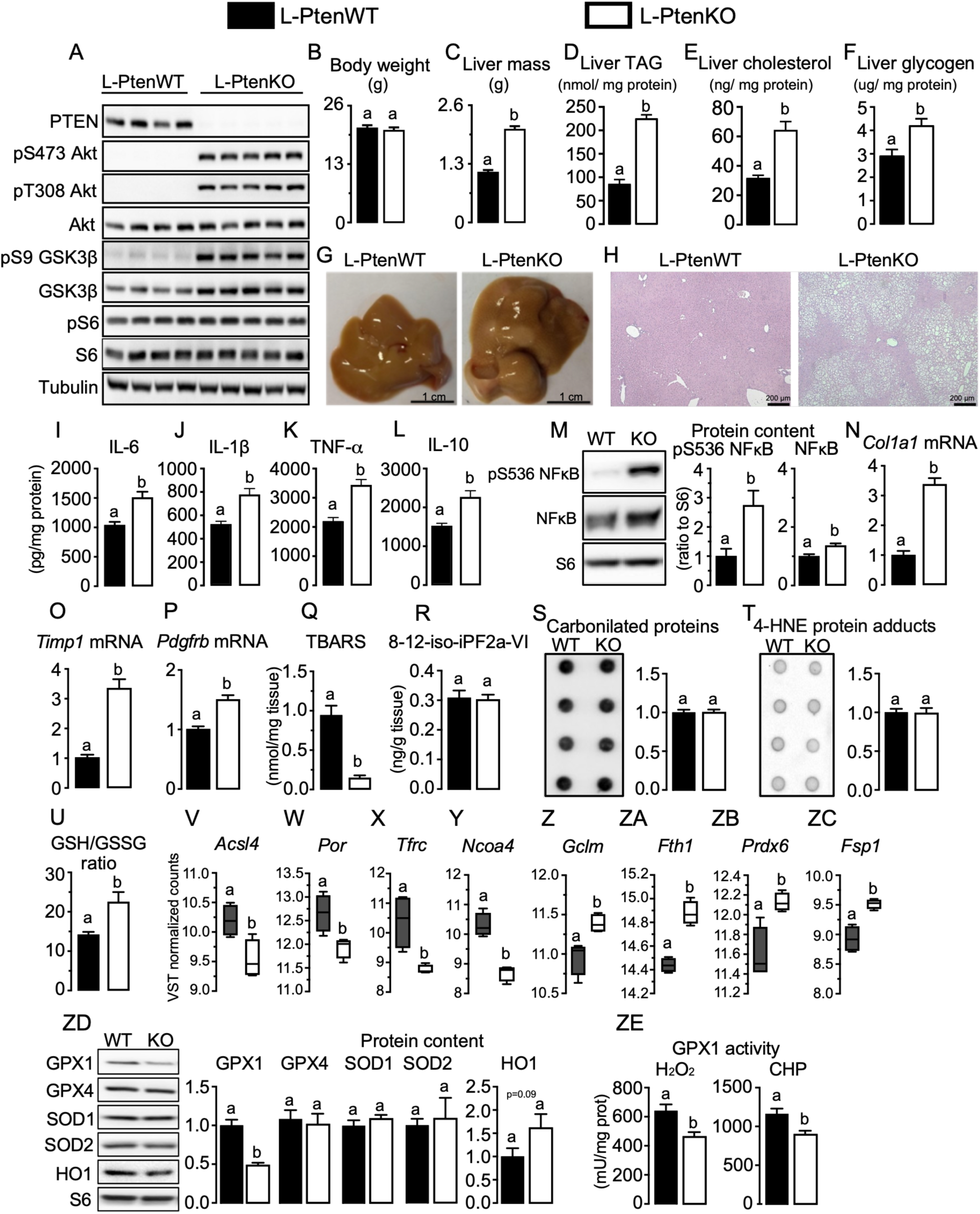
Constitutive PI3K-AKT activation in hepatocytes promotes steatohepatitis, hepatomegaly, inflammation and fibrosis, despite reducing lipid peroxidation and enhanced antioxidant capacity. (A) Representative Western blotting analysis of hepatic PTEN, phosphorylated AKT at Ser473 and Thr308, total AKT, phosphorylated GSK3β at Ser9, total GSK3β, phosphorylated S6, total S6, and tubulin. (B) Body weight, (C) liver mass, and hepatic (D) triacylglycerol (TAG), ® cholesterol, and (F) glycogen contents. (G) Representative macroscopic images of livers and (H) representative liver sections stained with hematoxylin and eosin (H&E). Hepatic (I–L) IL-6, IL-1β, TNF-α, and IL-10 contents and (M) NF-κB signaling, including phosphorylated NF-κB at Ser536 and total NF-κB protein abundance. Hepatic mRNA expression of (N) *Col1a1*, (O) *Timp1*, and (P) *Pdgfrb*, contents of (Q) thiobarbituric acid-reactive substances (TBARS), ® 8-12-isoprostane-iPF2a-VI, (S) carbonylated protein, (T) 4-hydroxynonenal (4-HNE) protein adducts, (U) reduced glutathione (GSH)/glutathione disulfide (GSSG) ratio. Hepatic mRNA expression of (V) *Acsl4*, (W) *Por*, (X) *Tfrc*, (Y) *Ncoa4*, (Z) *Gclm*, (ZA) *Fth1*, (ZB) *Prdx6*, (ZC) *Fsp1*, and (ZD) western blotting of glutathione peroxidase 1 and 4 (GPX1 and 4), superoxide dismutase 1 and 2 (SOD1) and (ZE) GPX1 activity. L-PtenWT, littermate control mice (*Pten* floxed); L-PtenKO, liver-specific *Pten* knockout mice (*Pten* floxed albumin-Cre+/−). Results are expressed as mean ± SEM. N = 5–7 mice per group. Student’s *t*-test was used to analyze the effect of *Pten* deletion (L-PtenWT vs. L-PtenKO). Means with different superscript letters differ significantly, *P* ≤ 0.05.

L-PtenKO mice also exhibited improved glucose tolerance and insulin sensitivity, reduced serum insulin levels (Supplementary Fig. 1A–C), increased inhibitory phosphorylation of FOXO1 at Ser256, reduced content of the gluconeogenic enzymes phosphoenolpyruvate carboxykinase (PEPCK) and glucose-6-phosphatase (G6Pase), and reduced glycemic response to pyruvate (Supplementary Fig. 1D–E). Interestingly, despite gluconeogenesis suppression, pyruvate carboxylase mRNA levels were increased in L-PtenKO liver (Supplementary Fig. 1F), suggesting a shift in pyruvate metabolism toward anaplerosis and replenishment of the tricarboxylic acid cycle intermediate oxaloacetate. *Ex vivo* liver slice assays revealed enhanced conversion of glucose into lactate, CO₂, glycogen, and triacylglycerol-glycerol in L-PtenKO at both fasting and refeeding. In contrast, glucose incorporation into triacylglycerol-fatty acids was enhanced only upon refeeding (Supplementary Fig. 1G–J).

Along with hepatomegaly, L-PtenKO displayed liver inflammation evidenced by the increased hepatic contents of IL-6 (Fig. 1I), IL-1β (Fig. 1J), TNFα (Fig. 1K) and IL-10 (Fig. 1L), and total and Ser536-phosphorylated NF-κB (Fig. 1M), and enhanced liver fibrosis evidenced by the increased mRNA contents of *Col1a1* (Fig. 1N), *Timp1* (Fig. 1O), and *Pdgfrb* (Fig. 1P). Surprisingly, despite pronounced inflammation, L-PtenKO livers displayed reduced lipid peroxidation as evidenced by the diminished TBARS content (Fig. 1Q), without, however, major alterations in oxidative damage evidenced by the unaltered liver content of 8-12-isoprostane F2a-VI (8-12-iso-iPF2a-VI, Fig. 1R), carbonylated proteins (Fig. 1S) and 4-hydroxynonenal (4-HNE) protein adducts (Fig. 1T). Interestingly, L-PtenKO livers exhibited increased GSH/GSSG ratio (Fig. 1U), and profound alterations in gene expression profile characterized by downregulation of genes favoring (*Acsl4*, *Por, Tfrc*, and *Ncoa4*, Fig. 1V-Y), in contrast to an upregulation of those protecting from lipid peroxidation and ferroptosis (*Gclm*, *Fth1*, *Prdx6* and *Fsp1*, Fig. 1Z-ZC). Along lipid peroxidation, L-PtenKO livers featured reduced glutathione peroxidase 1 (GPX1) content and activity, with unaltered GPX4, superoxide dismutase (SOD)1 and 2 and heme oxygenase 1 (HO1) contents (Fig. 1ZD and ZE). Thus, constitutive PI3K-AKT activation promotes steatohepatitis characterized by severe steatosis, inflammation, fibrosis, and hepatomegaly, which, surprisingly, is accompanied by reduced lipid peroxidation and enhanced antioxidant defense.

### Constitutive PI3K-AKT activation in hepatocytes remodels liver proteome by upregulating proteins involved in the opposite processes fatty acid *de novo* synthesis and oxidation

To further interrogate the mechanisms underlying L-PtenKO steatohepatitis, we performed a quantitative liver proteomic analysis (Fig. 2). Proteomics revealed extensive proteome remodeling in L-PtenKO livers, characterized by upregulation of 283 and downregulation of 270 proteins (Fig. 2A and Supplementary File 1). Functional enrichment analysis revealed, as expected, a marked upregulation of anabolic pathways, including glycogen metabolism, fatty acid biosynthesis (DNL), and glycolysis (Fig. 2B-C). In addition, proteins involved in triacylglycerol synthesis and lipolysis (CGI-58, MAGL, CES1D, CES1E), very-low density lipoprotein (VLDL) assembly, and glycerol 3-phosphate shuttle were also increased in L-PtenKO liver (Fig. 2C). Interestingly, along with anabolic processes, fatty acid β-oxidation was a significantly enriched pathway in L-PtenKO liver (Fig. 2B), featuring increased content of proteins involved in mitochondrial and peroxisomal fatty acid β-oxidation (Fig. 2C). Thus, constitutive PI3K-AKT activation in hepatocytes enhances both anabolic and catabolic arms of lipid metabolism, namely triacylglycerol synthesis and lipolysis, as well as fatty acid *de novo* synthesis (DNL) and β-oxidation.

**Figure 2.**
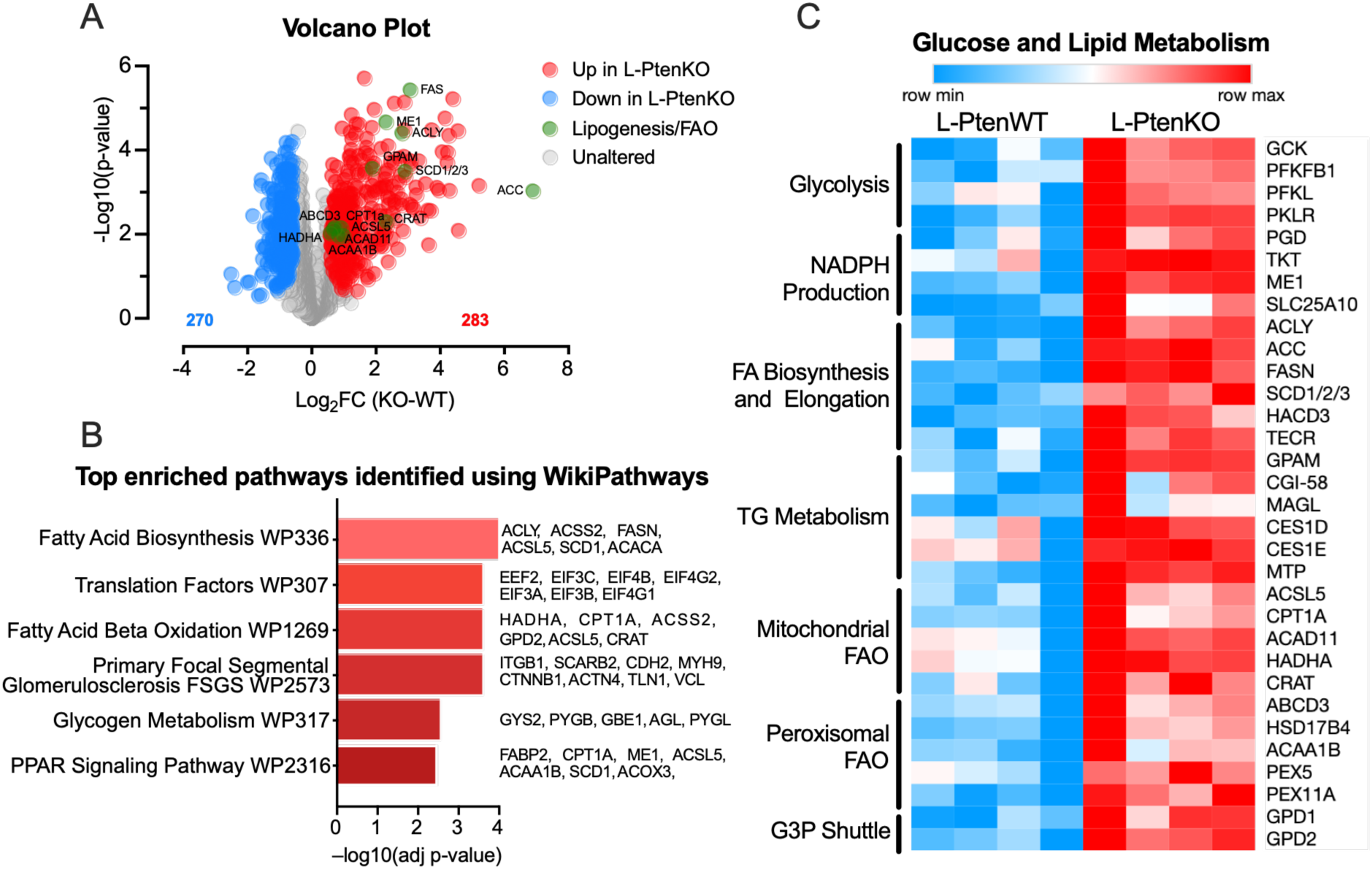
Constitutive PI3K-AKT activation in hepatocytes remodels liver proteome by upregulating proteins involved in the opposite processes fatty acid *de novo* synthesis and oxidation. (A) Volcano plot showing differentially abundant liver proteins between L-PtenWT and L-PtenKO mice identified by quantitative proteomic analysis. Proteins significantly increased and decreased in L-PtenKO livers are shown in blue and red, respectively, whereas proteins involved in de novo lipogenesis (DNL) and fatty acid oxidation (FAO) are highlighted in green. (B) Top enriched pathways among proteins significantly increased in L-PtenKO livers, identified using WikiPathways. (C) Heatmap showing the relative abundance of proteins involved in glucose and lipid metabolism in L-PtenWT and L-PtenKO livers. Protein abundance is represented by row-scaled Z-scores. L-PtenWT, littermate control mice (*Pten* floxed); L-PtenKO, liver-specific *Pten* knockout mice (*Pten* floxed albumin-Cre+/−). n = 4 mice per group. Differential protein abundance was determined using a two-sample Student’s *t*-test with permutation-based FDR correction in Perseus (FDR < 0.05, S0 = 0.5).

### Constitutive PI3K-AKT activation in hepatocytes enhances DNL and triacylglycerol synthesis and lipolysis (turnover)

Confirming and extending proteomic findings, hepatocyte *Pten* deletion markedly enhanced hepatic DNL, as evidenced by the increases in [¹⁴C]-acetate incorporation into fatty acids at both fasting and refeeding (Fig. 3A), and mRNA content of DNL-related enzymes *Acly, Acaca, Fasn,* and *Scd1*, and transcription factors sterol regulatory element-binding protein 1c (*Srebf1*) and carbohydrate-responsive element-binding protein β (*Chrebpβ*) (Fig. 3B). Protein content of ACC, FAS, and SCD1, as well as ACLY and FAS enzymatic activities (Fig. 3C–E) were also significantly increased in L-PtenKO liver. Along with DNL, hepatocyte *Pten* deletion enhanced triacylglycerol synthesis, as indicated by the increased incorporation of PA and LA into triacylglycerol (Fig. 3F–G) and total and NEM-resistant glycerol 3-phosphate acyltransferase (GPAT) activities at both fasting and refeeding (Fig. 3H). In contrast, NEM-sensitive GPAT activity was increased only in refed L-PtenKO mice (Fig. 3H). Importantly, hepatocyte Pten deletion upregulated both total and phosphorylated hormone-sensitive lipase (HSL) protein content (Fig. 3I), supporting enhanced triacylglycerol lipolysis and turnover in L-PtenKO liver. In line with these findings, RNA-seq revealed an upregulation of genes involved in triacylglycerol synthesis and lipolysis and lipid droplet remodeling, including *Mogat1, Fitm1, Plin3, Plin4, Cidea, Cidec, Abhd5,* and *Mgll* (Supplementary Fig. 2F-H). Altogether, metabolic studies confirm proteomic findings showing that constitutive activation of PI3K-AKT enhances DNL and triacylglycerol turnover in L-PtenKO mice livers.

**Figure 3.**
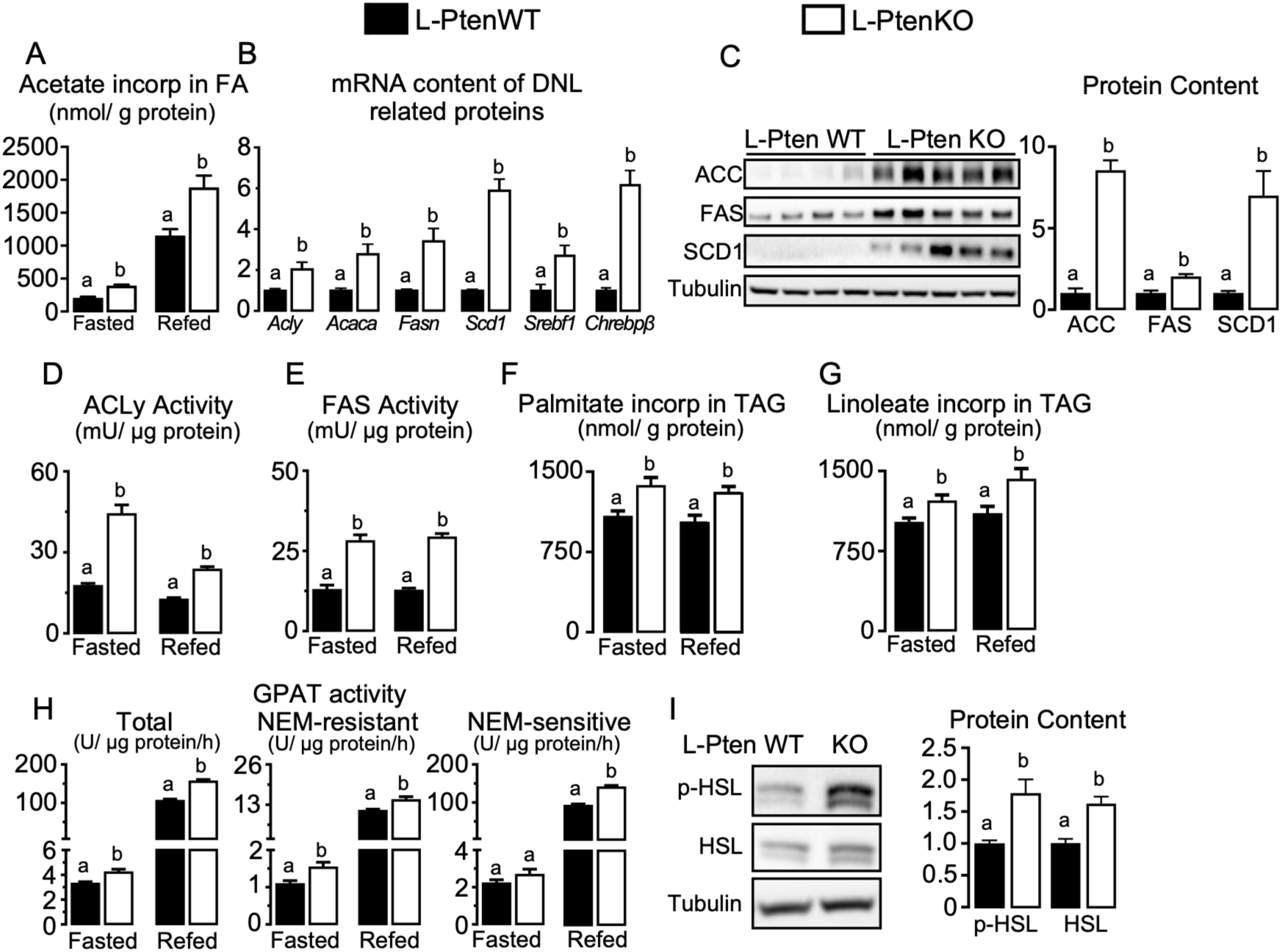
Constitutive PI3K-AKT activation in hepatocytes enhances DNL and triacylglycerol synthesis and lipolysis (turnover). (A) Hepatic [¹⁴C]-acetate incorporation into fatty acids in fasted and refed mice. (B) Hepatic mRNA expression of de novo lipogenesis (DNL)-related genes, including *Acly, Acaca, Fas, Scd1, Srebf1,* and *Chrebp*. (C) Hepatic protein abundance of ACC, FAS, and SCD1. (D–E) Hepatic ACLY and FAS enzymatic activities, respectively. (F–G) Ex vivo incorporation of palmitate and linoleate into triacylglycerol (TAG), respectively. (H) Total, N-ethylmaleimide (NEM)-resistant, and NEM-sensitive glycerol-3-phosphate acyltransferase (GPAT) activities. (I) Hepatic total HSL and phosphorylated HSL (Ser660) protein abundance. L-PtenWT, littermate control mice (*Pten* floxed); L-PtenKO, liver-specific *Pten* knockout mice (*Pten* floxed albumin-Cre+/−). Measurements were performed in fasted and refed 8-week-old mice. Results are expressed as mean ± SEM. n = 6–8 mice per group. Student’s *t*-test was used to analyze the effect of *Pten* deletion (L-PtenWT vs. L-PtenKO) within each nutritional state. Means with different superscript letters are significantly different, *P* ≤ 0.05.

### Constitutive PI3K-AKT activation in hepatocytes increases lipid species containing DNL-derived fatty acids and depletes those containing essential fatty acids

Next, we investigated the impact of constitutive PI3K-AKT activation on liver lipidome. Untargeted liver lipidomics identified 307 lipid species across 16 lipid subclasses, primarily phospholipids (129 species), neutral lipids (113 species), and sphingolipids (26 species) (Supplementary File 2). Principal component analysis (PCA) revealed a clear separation between L-PtenWT and L-PtenKO (Fig. 4A) livers, indicating extensive lipidome remodeling (Fig. 4A) mainly characterized by increased abundance of lipid species containing the DNL-related fatty acids PA (16:0), POA (16:1), SA (18:0) and OA (18:1), accompanied by a marked reduction in species containing the essential fatty acids LA (18:2) and ALA (18:3) (Fig. 4B). This pattern was consistently found in multiple lipid classes including phosphatidylcholine (PC), phosphatidylethanolamine (PE), and phosphatidylglycerol (PG) (Fig. 4C–E), as well as cholesteryl esters (CE), triacylglycerol (TG), diacylglycerol (DG), and free fatty acids (Fig. 4H–K). Exceptions to this pattern were the reduction in phosphatidylinositol (PI) species containing essential fatty acids without changes in those containing DNL-derived fatty acids (Fig. 4F), and the broad cardiolipin reduction independently of fatty acid composition (Fig. 4G). Lipid species containing PUFAs (AA 20:4 n-6, EPA 20:5 n-3, DHA 22:6 n-3, docosatetraenoic acid 22:4 and DPA 22:5) showed a heterogeneous behavior, with some species being upregulated (PC, PG, and DG), while others (PE, cardiolipin, CE, and TG) were downregulated in L-PtenKO liver. Of note, L-PtenKO marked depletion in liver LA- and ALA-containing lipids was associated with a broad reduction in content of their oxygenated derivatives (Fig. 4L), excluding a role of LA- and ALA-derived oxylipin generation as a major determinant of essential fatty acid depletion in these mice. Next, GC-FAME analysis of fasted and refed mice confirmed the enrichment in liver DNL-derived fatty acids and the essential fatty acids depeltion in L-PtenKO livers across feeding states (Supplementary Fig. 3A–B and Tables 4 and 5). L-PtenKO mice also showed a lower hepatic AA (20:4n6) content than L-PtenWT at fasting, but not refeeding, and a lower docosahexaenoic acid (DHA, 22:6n3) content than L-PtenWT mice at both conditions (Supplementary Fig. 3A–B, Supplementary Tables 3 and 4). Furthermore, the difference between the contents of LA and its elongated product di-homo-γ-linolenic acid (DGLA, 20:3n-6) was similar at fasting and refeeding (Supplementary Fig. 3C-D) and, among the enzymes involved in LA conversion to AA, fatty acid desaturase 2 (*Fads2*), the first and rate-limiting, had its mRNA content downregulated, while that of fatty acid elongase 5 (*Elovl5*) and *Fads1* were upregulated in L-PtenKO liver (Supplementary Fig. 3E). Thus, constitutive PI3K-AKT and DNL activation promotes a robust liver lipidome remodeling characterized by enrichment of lipid species containing DNL-derived fatty acids, as well as by a marked depletion of lipid species composed of essential fatty acids, a lipid signature that is not affected by feeding status and is not due to LA elongation and desaturation to AA or conversion to oxidized derivatives.

**Figure 4.**
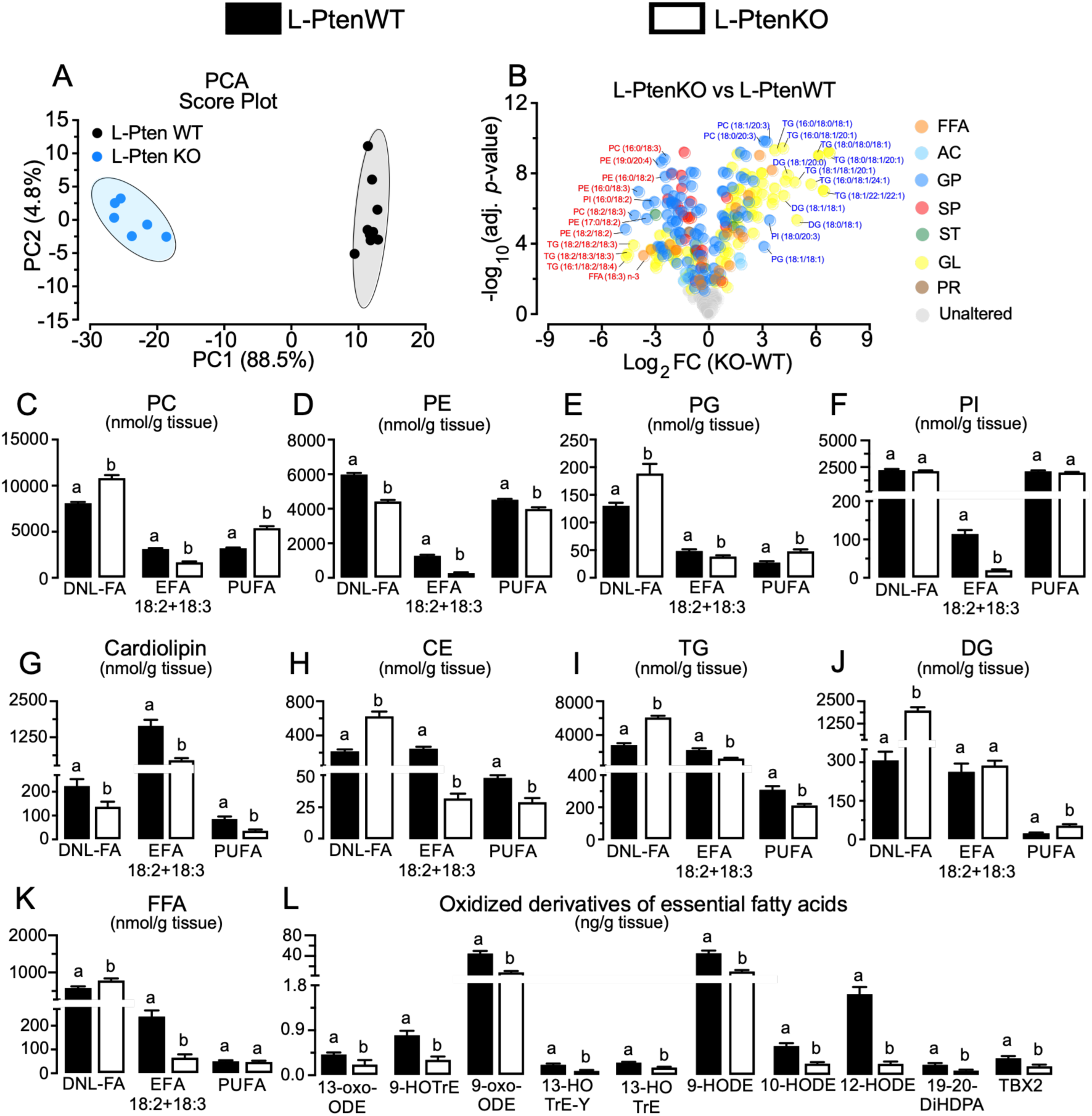
Constitutive PI3K-AKT activation in hepatocytes increases lipid species containing DNL-derived fatty acids and depletes those containing essential fatty acids. **(A)** Principal component analysis (PCA) of hepatic lipidomic profiles. **(B)** Volcano plot of differentially abundant lipid species between L-PtenKO and L-PtenWT livers, with log₂ fold change plotted against −log₁₀ adjusted *P* value. Lipid species are colored according to lipid class, including free fatty acids (FFA), acylcarnitines (AC), glycerophospholipids (GP), sphingolipids (SP), sterol lipids (ST), glycerolipids (GL), and prenol lipids (PR). **(C–K)** Hepatic levels of phosphatidylcholine (PC), phosphatidylethanolamine (PE), phosphatidylglycerol (PG), phosphatidylinositol (PI), cardiolipin (CL), cholesteryl esters (CE), triacylglycerol (TG), diacylglycerol (DG), and free fatty acids (FFA), respectively, grouped according to fatty acid composition as DNL-derived fatty acids (DNL-FA), essential fatty acids (EFA; 18:2 and 18:3), and polyunsaturated fatty acids (PUFA). **(L)** Hepatic levels of oxidized derivatives of essential fatty acids. Lipidomic analysis was performed in 8-week-old L-PtenWT and L-PtenKO mice. L-PtenWT, littermate control mice (*Pten* floxed); L-PtenKO, liver-specific *Pten* knockout mice (*Pten* floxed albumin-Cre+/−). Results are expressed as mean ± SEM. *n* = 7–9 mice per group. Statistical significance for individual lipid species was determined using Student’s *t*-test followed by false discovery rate (FDR) correction and a fold-change threshold of >1.0. Means with different superscript letters are significantly different, *P* ≤ 0.05.

### Constitutive PI3K–AKT activation in hepatocytes increases fatty acid oxidation and mitochondrial respiratory capacity while altering mitochondrial dynamics

Given the enrichment of mitochondrial and peroxisomal proteins revealed by proteomics, we next investigated whether increased β-oxidation could contribute to essential fatty acid depletion. Indeed, L-PtenKO steatohepatitis was associated with a marked increase in both fatty acid β-oxidation and carbon flux through the tricarboxylic acid cycle, as evidenced by the increased [¹⁴C]-palmitate, [¹⁴C]-linoleate and [¹⁴C]-acetate conversions into ¹⁴CO₂, respectively, at both fasting and refeeding, with more pronounced effects during fasting (Fig. 5A–C). Interestingly, L-PtenKO livers also exhibited increased citrate synthase activity (Fig. 5D), a surrogate marker of mitochondrial mass, along with increased coupled (state 3) and maximal uncoupled (state 3U) mitochondrial respiration using either succinate or palmitoyl- or linoleoyl-carnitine plus malate as substrates (Fig. 5E–G), altogether indicating enhanced mitochondrial β-oxidation, electron transport activity, and oxidative capacity. L-PtenKO livers also exhibited increased non-phosphorylating respiration (state 4) in the presence of succinate (Fig. 5E), suggesting enhanced proton leak, which may be secondary to the elevated electron flux in the respiratory chain. Finally, there were no differences in the respiratory control ratio (RCR) between genotypes (Fig. 5H), indicating a preservation of mitochondrial coupling efficiency, despite the increased respiration.

**Figure 5.**
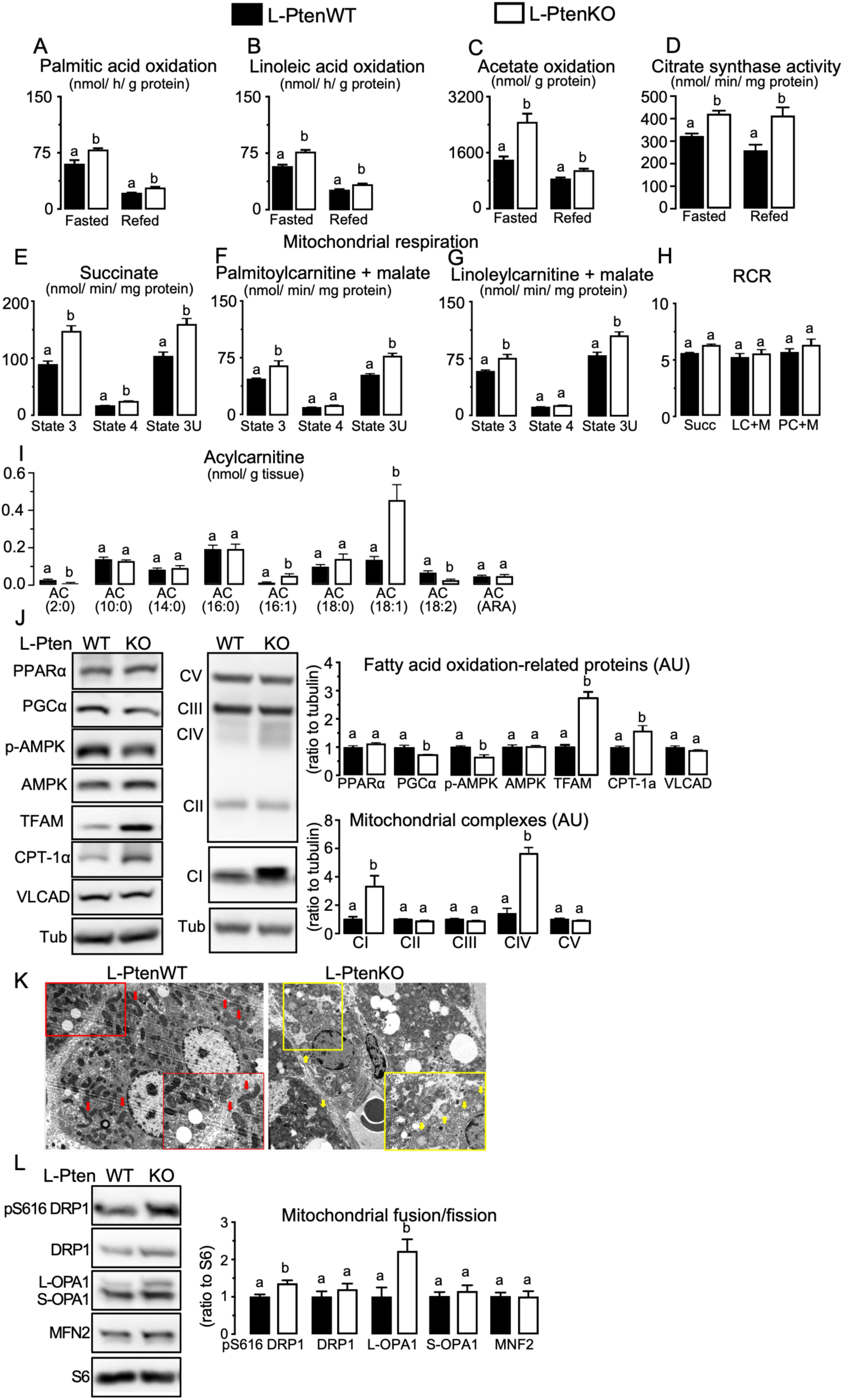
Constitutive PI3K–AKT activation in hepatocytes increases fatty acid oxidation and mitochondrial respiratory capacity while altering mitochondrial dynamics. **(A–C)** Hepatic oxidation of [1-¹⁴C]palmitic acid, [1-¹⁴C]linoleic acid, and [1-¹⁴C]acetate, respectively, in fasted and refed mice. **(D)** Hepatic citrate synthase activity in fasted and refed mice. **(E–G)** Oxygen consumption rates in isolated liver mitochondria measured at State 3 (1 mM ADP), State 4 (0.5 μg/mL oligomycin), and State 3U (CCCP) using (E) succinate (5 mM) plus rotenone (1 μM), (F) palmitoylcarnitine (25 μM) plus malate (2 mM), and (G) linoleylcarnitine (25 μM) plus malate (2 mM) as respiratory substrates. **(H)** Respiratory control ratio (RCR), calculated as the State 3/State 4 respiration ratio, using succinate (Succ), linoleylcarnitine plus malate (LC+M), or palmitoylcarnitine plus malate (PC+M). **(I)** Hepatic acylcarnitine levels. **(J)** Hepatic protein abundance of PPARα, PGC-1α, phosphorylated AMPK, total AMPK, TFAM, CPT1α, and VLCAD, and mitochondrial respiratory chain complexes I–V. Tubulin was used as a loading control. **(K)** Representative transmission electron microscopy images of liver mitochondria from L-PtenWT and L-PtenKO mice. Red arrows indicate elongated mitochondria predominantly observed in L-PtenWT livers, whereas yellow arrows indicate more rounded mitochondria with disorganized mitochondrial matrix predominantly observed in L-PtenKO livers. **(L)** Hepatic protein abundance of phosphorylated DRP1 at Ser616, total DRP1, long and short OPA1 isoforms, and MFN2 in fasted and refed mice. L-PtenWT, littermate control mice (*Pten* floxed); L-PtenKO, liver-specific *Pten* knockout mice (*Pten* floxed albumin-Cre+/−). Results are expressed as mean ± SEM. Student’s *t*-test was used to analyze the effect of *Pten* deletion (L-PtenWT vs. L-PtenKO) within each nutritional state. Means with different superscript letters are significantly different, *P* ≤ 0.05.

Consistently with a preferential LA oxidation, acylcarnitine profiling revealed increased content of species containing DNL-associated fatty acids, particularly C16:1 and C18:1, whereas that containing the essential fatty acid LA namely linoleoylcarnitine (C18:2) was reduced in L-PtenKO livers (Fig. 5I). L-PtenKO mice also exhibited increased hepatic CPT1α, mitochondrial transcription factor A (TFAM), and mitochondrial respiratory complexes I and IV (Fig. 4J), unaltered PPARα, AMP-activated protein kinase (AMPK), and very long-chain acyl CoA dehydrogenase (VLCAD) and reduced PPAR coactivator 1α (PGC1α) and phosphorylated AMPK protein contents (Fig. 5J). Despite unaltered PPARα, but consistently with reduced PGC1α content, RNA-seq analysis revealed a downregulation of the canonical PPARα target genes *Cyp4a10, Cyp4a14, Acot1, Ehhadh, Hadha,* and *Hadhb,* along with unaltered *Cpt1a* (Supplementary Fig. S2). Thus, constitutive PI3K-AKT and DNL activation enhances fatty acid β-oxidation, carbon flux in the tricarboxylic cycle flux, mitochondrial mass, respiration and oxidative capacity along a depletion of acylcarnitine-containing LA. Mechanistically, such elicitation of mitochondrial oxidative machinery may be driven in part by TFAM and seems independent of PGC1α and PPARα.

One puzzling finding was that constitutive PI3K-AKT activation enhanced mitochondrial mass and respiration despite reducing total and LA-containing cardiolipins (CL) (Supplementary Fig. S4A), unique phospholipids of the inner mitochondrial membrane that contribute to cristae organization, respiratory complexes stabilization and activation, and electron flux [62,63]. Indeed, as revealed by RNAseq, L-PtenKO cardiolipin reduction in liver was associated with diminished mRNA content of cardiolipin synthase 1 (*Crls1*), which catalyzes *de novo* cardiolipin synthesis, and patatin-like domain 8, phospholipase A2 (*Pnpla8*), which initiates cardiolipin de-acylation and remodeling, while mRNA content of tafazzin (*Taz*), which re-acylates monolysocardiolipin during cardiolipin remodeling, was significantly increased (Supplementary Fig. 4B-C). Consistently with the reduced *Crls1*, phosphatidylglycerol (PG), the immediate substrate for cardiolipin synthesis, was increased in L-PtenKO livers (Fig. 4E). Notably, in agreement with cardiolipin structural role determining mitochondria inner membrane curvature, transmission electron microscopy revealed poorly developed and disorganized mitochondrial cristae in L-PtenKO liver (Fig. 5K), which were associated with increased contents of long optic atrophy 1 (L-OPA1), that maintains mitochondrial cristae morphology, and dynamin related GTPase (DRP1) phosphorylated at Ser616, involved in mitochondrial fission (Fig. 5L). Liver contents of mitofusin 2 (MFN2), total DRP1, and short-

OPA1 (S-OPA1), which also act on mitochondrial remodeling, were not altered (Fig. 5L). Proteomic analysis further revealed significant increases of proteins involved in mitochondrial maintenance namely lon peptidase 1 (LONP1), that maintains protein quality control in mitochondrial matrix; inner membrane mitochondrial protein (IMMT), that supports cristae formation and inner mitochondrial membrane organization; and AFG3 Like Matrix AAA Peptidase Subunit 2 (AFG3L2), involved in mitochondrial proteostasis, while contents of caseinolytic mitochondrial matrix peptidase proteolytic subunit (CLPP), involved in matrix proteostasis, metataxin 1 (MTX1), that carries mitochondrion organization and protein transport, and apolipoprotein O (APOO), that supports mitochondrial cristae junction integrity and architecture, were reduced in L-PtenKO livers (Supplementary Fig. 4E). Altogether, the enhanced respiration in L-PtenKO mitochondria was associated with increased L-OPA1, DRP1, LONP1 and IMMT involved in cristae organization and mitochondrial proteostasis, despite reduced cardiolipin content and altered mitochondrial cristae architecture.

### Dietary LA supplementation restores hepatic LA depletion and partially attenuates inflammation induced by constitutive PI3K-AKT activation in hepatocytes

Considering LA and AA roles in lipid peroxidation and inflammation [59], we next investigated whether dietary LA supplementation could not only abrogate liver LA depletion, but also impact lipid peroxidation, inflammation and SLD progression in L-PtenKO mice. For this, L-PtenWT and L-PtenKO mice were fed either LF, HF, or HFLA diets containing increasing amounts of LA for 4 weeks. As depicted in Figure 6, LF-fed L-PtenWT mice had higher liver LA content than LF-fed L-PtenKO. HF intake similarly increased liver LA content in both L-PtenWT and L-PtenKO, maintaining LA content higher in the former. HFLA intake, on the other hand, further increased hepatic LA content in L-PtenKO, but not L-PtenWT, abolishing the difference between genotypes (Fig. 6A). In contrast to LA, hepatic AA (20:4n-6) levels did not differ between L-PtenWT and L-PtenKO mice under any dietary condition, although both HF and HFLA feeding modestly increased AA levels in L-PtenKO mice (Fig. 6B). Conversely, hepatic DNL-derived fatty acid content was mildly reduced in L-PtenKO by HFLA, but not HF intake, when compared to LF-fed L-PtenKO, but levels remained higher in L-PtenKO than L-PtenWT in all diets (Fig. 6C). Along liver LA replenishment, HFLA, but not HF intake, moderately reduced L-PtenKO liver mass in comparison to LF-fed L-PtenKO (Fig. 6D). In spite of this reduction, liver mass was significantly higher in L-PtenKO than L-PtenWT in all diets investigated. Liver triacylglycerol content and steatosis, which were higher in L-PtenKO than L-PtenWT in all diets, were not affected by either HF or HFLA intake (Fig. 6E-F). *Ex vivo* metabolic tracing showed reduced acetate incorporation into fatty acids (DNL) in L-PtenKO mice fed with both HF and HFLA (Fig. 6G). Neither HF nor HFLA intake affected liver TBARS content, which was lower in L-PtenKO than L-PtenWT regardless of diet (Fig. 6H). In contrast to TBARS, hepatic levels of the proinflammatory cytokines IL-6 and IL-1β were consistently higher in L-PtenKO than in L-PtenWT, independently of diet (Fig. 6I-J). Importantly, hepatic content of TNF-α and IL-10 was higher in L-PtenKO than L-PtenWT fed with either LF or HF, while HFLA intake reduced liver TNF-α and IL-10 content in L-PtenKO, abolishing the difference between genotypes (Figure 6K-L). In contrast to inflammation, liver fibrosis was not significantly affected by dietary LA supplementation, as evaluated by *Col1a1* and *Timp1* mRNA levels (Figure 6M-N). Altogether, dietary supplementation and restoration of liver LA content in L-PtenKO mice reduced liver inflammation without affecting fibrosis.

**Figure 6.**
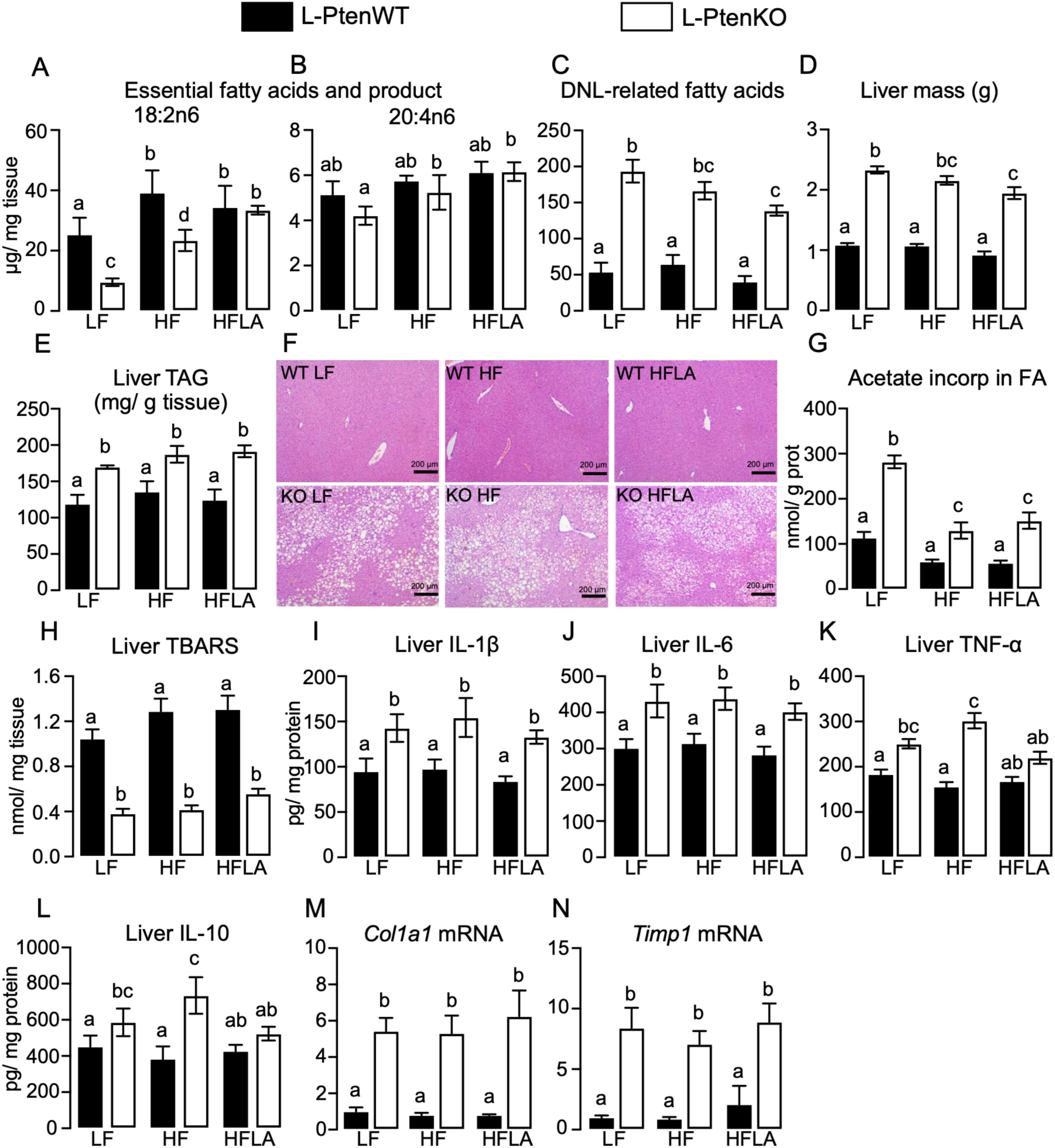
Dietary LA supplementation restores hepatic LA depletion and partially attenuates inflammation induced by constitutive PI3K-AKT activation in hepatocytes. **(A)** Hepatic linoleic acid (LA; 18:2n-6) content, **(B)** hepatic arachidonic acid (AA; 20:4n-6) content, and **(C)** hepatic DNL-derived fatty acid content. **(D)** Liver mass and **(E)** hepatic triacylglycerol (TAG) content. **(F)** Representative liver sections stained with hematoxylin and eosin (H&E). **(G)** Hepatic [¹⁴C]-acetate incorporation into fatty acids. **(H)** Hepatic thiobarbituric acid-reactive substances (TBARS) content. Hepatic **(I)** IL-1β, **(J)** IL-6, **(K)** TNF-α, and **(L)** IL-10 contents. Hepatic mRNA expression of (M) *Col1a1* and (N) *Timp1*. L-PtenWT and L-PtenKO mice were fed a low-fat diet (LF; 14.4 g/kg LA), a high-fat diet (HF; 41 g/kg LA), or a high-fat diet supplemented with LA (HFLA; 77.3 g/kg LA). Mice were fasted for 10 h and refed for 3 h before tissue collection. Results are expressed as mean ± SEM. *n* = 5–8 mice per group. Statistical differences were analyzed by two-way ANOVA followed by Tukey’s post hoc test. Means with different superscript letters are significantly different, *P* ≤ 0.05.

### Pharmacological ACC inhibition reduces DNL and steatosis, but further depletes essential fatty acids in liver of mice bearing constitutive PI3K-AKT activation in hepatocytes

We next interrogated the relative contribution of DNL to L-PtenKO liver steatohepatitis by pharmacologically inhibiting acetyl-CoA carboxylase (ACC) (Fig. 7). In addition to reducing DNL, pharmacological ACC inhibition is expected to alleviate CPT1α allosteric inhibition by malonyl-CoA, enhancing fatty acid β-oxidation. ACC inhibition did not alter the hepatomegaly, namely, the increase in liver mass induced by hepatocyte Pten deletion (Fig. 7A), despite markedly reducing liver triacylglycerol content, acetate incorporation in fatty acids, steatosis and content of DNL-derived fatty acids (Fig. 7B–E). Interestingly, ACC inhibition did not affect liver LA in L-PtenWT, but further reduced its content in L-PtenKO mice (Fig. 7F). Hepatic AA levels, on the other hand, were reduced by ACC inhibition in both L-PtenWT and L-PtenKO mice (Fig. 7G). Similarly to LA, ACC inhibition tended to reduce ALA content in L-PtenKO (Fig. 7H), but significantly reduced the content of its product DHA in both L-PtenWT and L-PtenKO mice (Fig. 7I). ACC inhibition did not affect liver contents of TBARs (Fig. 7J) and of phosphorylated and total AKT2 and phosphorylated ACC, but further increased contents of ACC and FAS in L-PtenKO mice (Fig. 7K). Furthermore, although NF-κB phosphorylation was unaltered, total NF-κB protein abundance was markedly reduced following ACC inhibition (Fig. 7K). Corroborating these findings, ACC inhibition significantly reduced hepatic inflammation and fibrosis in L-PtenKO mice, as evidenced by the reduced liver content of IL-6, TNFα, and IL10, but not IL-1β (Fig. 7L-O) and mRNA content of *Col1a1* and *Timp1* (Fig. 7P-Q), respectively. Altogether, these findings indicate that pharmacological ACC and DNL inhibition enhances essential fatty acid depletion and attenuate steatosis, inflammation, and fibrosis in L-PtenKO liver.

**Figure 7.**
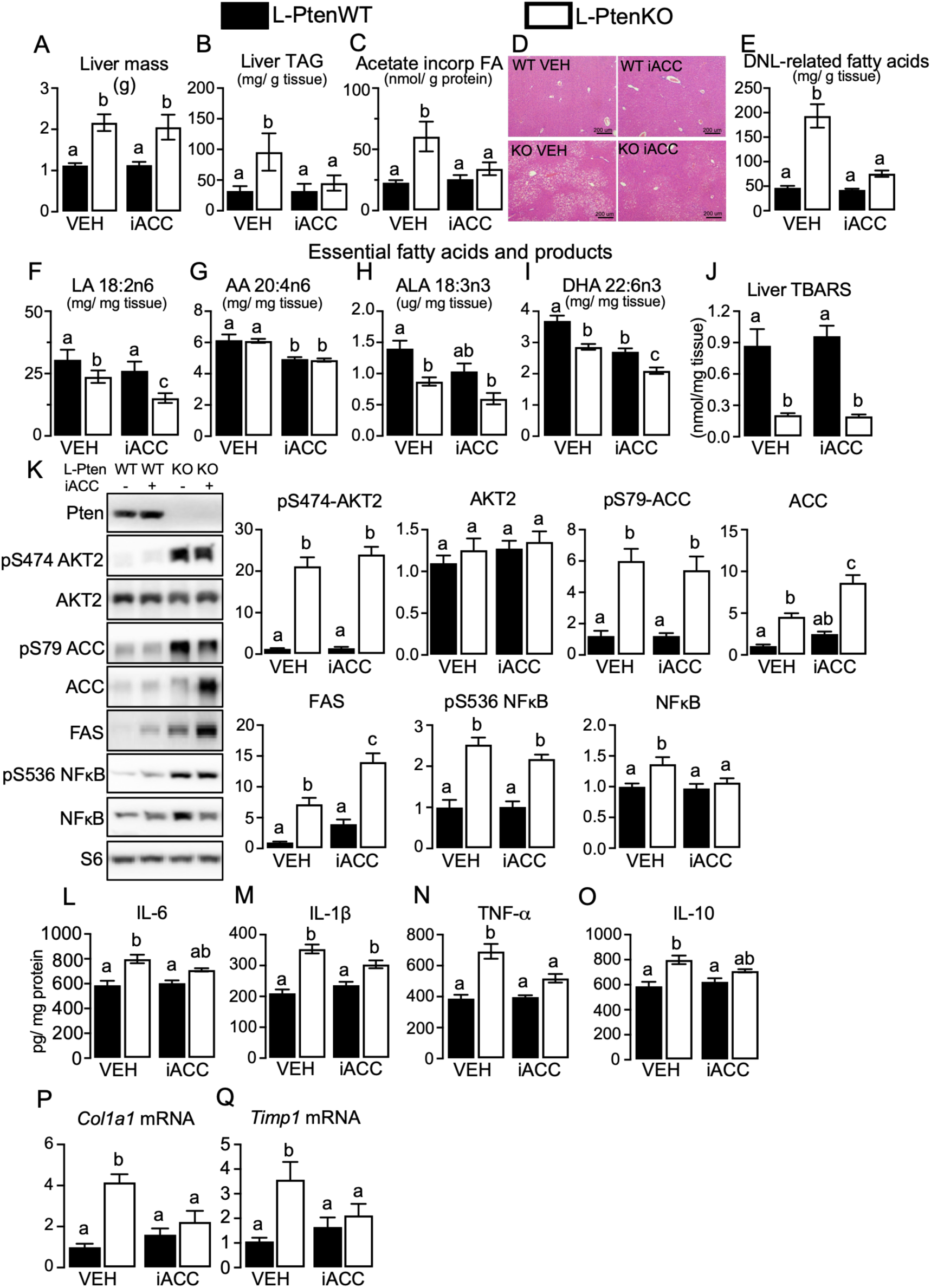
Pharmacological ACC inhibition reduces DNL and steatosis, but further depletes essential fatty acids in liver of mice bearing constitutive PI3K-AKT activation in hepatocytes. (A) Liver mass, (B) hepatic triacylglycerol (TAG) content, (C) hepatic [¹⁴C]-acetate incorporation into fatty acids, (D) representative H&E-stained liver sections, and (E) hepatic DNL-derived fatty acid content. (F–I) Hepatic levels of linoleic acid (LA; 18:2n-6), arachidonic acid (AA; 20:4n-6), α-linolenic acid (ALA; 18:3n-3), and docosahexaenoic acid (DHA; 22:6n-3), respectively. (J) Hepatic thiobarbituric acid-reactive substances (TBARS) content. (K) Hepatic protein abundance of PTEN, phosphorylated AKT2 at Ser474, total AKT2, phosphorylated ACC at Ser79, total ACC, FAS, phosphorylated NF-κB at Ser536, and total NF-κB. S6 was used as a loading control. (L–O) Hepatic IL-6, IL-1β, TNF-α, and IL-10 contents, respectively. (P–Q) Hepatic *Col1a1* and *Timp1* mRNA expression, respectively. L-PtenWT and L-PtenKO mice were treated with vehicle (VEH; 1% DMSO in 0.2% carboxymethylcellulose) or the ACC inhibitor ND-630 (iACC; 20 mg/kg/day, i.p.). Results are expressed as mean ± SEM. *n* = 5–7 mice per group. Statistical differences were analyzed by two-way ANOVA followed by Tukey’s post hoc test. Means with different superscript letters are significantly different, *P* ≤ 0.05.

## 4. Discussion

Constitutive PI3K-AKT and DNL activation in hepatocytes promoted steatohepatitis characterized by severe steatosis, hepatocyte ballooning, inflammation, fibrosis and hepatomegaly, which was unexpectedly associated with reduced lipid peroxidation, increased GSH content and enhanced rates of triacylglycerol turnover, fatty acid β-oxidation, tricarboxylic acid cycle flux, and mitochondrial respiration, altogether promoting a robust lipidome remodeling mainly defined by enrichment of DNL-derived fatty acids in detriment of a broad depletion of essential fatty acids (LA and ALA) and cardiolipins. Dietary restoration of liver LA content mildly attenuated liver inflammation, but not steatosis and fibrosis, while pharmacological ACC and DNL inhibition robustly mitigated L-PtenKO mice hepatic steatosis, inflammation and fibrosis, at the expense of further enhancing hepatic essential LA depletion, such findings that support DNL implication as a major culprit of steatohepatitis induced by constitutive PI3K-AKT activation.

Although chronic oxidative stress is a hallmark of SLD, despite robust lipid accumulation, steatohepatitis induced by constitutive PI3K-AKT activation occurred along reduced oxidative stress and lipid peroxidation, such phenotypes that may be attributed to a broad recruitment of antioxidant defense system characterized by reduced mRNA content of genes involved in reactive oxygen species generation, PUFA peroxidation and iron metabolism (*Acsl4, Por, Tfrc* and *Ncoa4*) along an upregulation of genes involved in antioxidant response, GSH synthesis and protection from iron-driven lipid peroxidation (*Gclm*, *Fth1*, *Prdx6*, and *Fsp1*). Corroborating these findings, cells bearing constitutive PI3K-AKT signaling showed enhanced resistance to oxidative stress and ferroptosis, a response that was mechanistically attributed to either mitochondrial fitness [64] or increased SREBP1-SCD1-mediated synthesis of monounsaturated fatty acids [65]. Supporting the former, constitutive PI3K-AKT activation was indeed associated with increased mitochondrial mass and respiration, while in contrast to the latter, pharmacological ACC and DNL inhibition significantly reduced liver content of DNL-related fatty acids without affecting liver lipid peroxidation. This redox phenotype found in 8 weeks old mice contrasts with the marked lipid peroxidation and nuclear factor erythroid 2-related factor 2 (Nrf2) activation reported in 45–55 weeks old L-PtenKO mice [66], in which the liver is severely overtaken by tumors, suggesting an age-dependent transition towards a pro-oxidant redox status upon persistence of metabolic stress and progression of SLD induced by PTEN deficiency [67].

In addition to the reduced lipid peroxidation, steatohepatitis induced by constitutive PI3K-AKT activation was associated with activation of important lipid-related futile metabolic cycles as the result of simultaneous activation of the opposite processes, triacylglycerol synthesis and lipolysis, as well as fatty acid *de novo* synthesis (DNL) and β-oxidation. Specifically to the former, simultaneously to the enhanced glucose conversion in glycerol 3-phosphate, GPAT activity and fatty acid esterification in triacylglycerol, constitutive PI3K-AKT activation upregulated proteins involved in triacylglycerol lipolysis including the adipose triglyceride lipase (ATGL) coactivator CGI-58, phosphorylated HSL, monoacylglycerol lipase (MAGL) [68] along with the endoplasmic reticulum carboxylesterase CES1D, a neutral lipid hydrolase that enhances triacylglycerol turnover by activating the transcriptional factors PPARγ and SREBP1c [69]. Interestingly, PPARγ content and activity are markedly upregulated in SLD induced by hepatocyte *Pten* deficiency, wherein this nuclear receptor is responsible for approximately 50% of liver hypertrophy [70]. Considering that PPARγ activation was previously shown to enhance both arms of the triacylglycerol-fatty acid cycle in adipocytes [71], it emerges as a likely candidate to mediate the activation of this metabolic cycle in L-Pten KO liver, a hypothesis that will be tested in future studies.

Although simultaneous activation of DNL and β-oxidation seems paradoxical considering CPT1α inhibition by malonyl-CoA, activation of this cycle has been previously recognized in brown adipose tissue upon cold-induced thermogenesis [72,73] and in the livers of mice bearing SLD induced by severe lipoatrophy [29], and HCC-bearing patients [28]. Possible mechanisms allowing bypass of CPT1α allosteric inhibition under these conditions may include a rapid malonyl-CoA utilization via enhanced FAS activity, increased mitochondrial malonyl-CoA degradation by malonyl-CoA decarboxylase (MCD), as well as the distinct subcellular malonyl-CoA pools produced by ACC1 and ACC2, among others [72,74]. Mechanistically, the increase in both mRNA and protein contents of ACLY, ACC, FAS and SCD1 found here indicates that DNL activation in L-PtenKO is most likely induced at the transcriptional level by SREBP1 and ChREBP, while the mechanisms driving the upregulation in β-oxidation are still not completely defined. In this context, despite previous evidence [30], our findings showing unaltered liver PPARα mRNA and protein content in L-PtenKO, along a marked reduction in liver mRNA levels of several canonical PPARα target genes, which may be partly explained by decreased levels of the PPARα coactivator PGC-1α, argues against an involvement of PPARα as mediator of these phenotypes. Additionally, CPT1α, the rate-limiting enzyme of β-oxidation, which is transcriptionally induced by PPARα was upregulated in L-PtenKO liver at the protein, but not mRNA level, indicating a post-transcriptional regulation. Aside its regulation, β-oxidation activation in L-PtenKO livers was associated with higher carbon flux in the tricarboxylic acid cycle and increased mitochondrial mass, contents of respiratory complexes I and IV and respiration. Interestingly, enhanced flux in the tricarboxylic acid cycle occurred along with increased expression of pyruvate carboxylase, which may sustain the elevated cycle activity by catalyzing the anaplerotic production of its intermediate, oxaloacetate. Mechanistically, L-PtenKO liver enhanced mitochondrial mass and respiration, which energetically support these energy-expending metabolic cycles, were associated with increased contents of TFAM, which enhances transcription of mitochondrial genes coding respiratory complexes, as well as the RNA-binding protein clustered mitochondria homolog (CLUH), which was previously shown to enhance the translation of mRNAs encoding mitochondrial proteins <u>[75]</u>. This post-transcriptional mechanism may explain, at least in part, the increased abundance of proteins involved in mitochondrial β-oxidation, including CPT1α, without a corresponding increase in mRNA content.

Elicitation of metabolic cycles, as seen in L-PtenKO livers, serves different purposes, including increasing energy expenditure, enhancing sensitivity to hormonal control, and accelerating the remodeling of fatty acids composing lipid species, among others [76]. Supporting the latter, lipidome analysis of L-PtenKO liver revealed robust lipid remodeling characterized by not only enrichment of DNL-derived fatty acids, as expected considering the constitutive DNL activation, but also broad depletion of the essential fatty acids LA and ALA and cardiolipins. Indeed, considering that liver LA depletion occurred independently of the feeding status and was not accompanied by a proportional equimolar increase in LA-derived products including the PUFAs DGLA and AA or the oxidized lipid derivatives HODEs and oxo-ODEs, such depletion is most likely caused by LA preferential β-oxidation. Supporting this notion, L-PtenKO livers display reduced LA-derived linoleoylcarnitine, which may indicate preferential oxidation, and pharmacological ACC inhibition, which enhances β-oxidation by reducing malonyl-CoA-mediated inhibition of CPT1α, further reduced hepatic LA levels in L-PtenKO livers. Furthermore, previous *in vivo* and *in vitro* studies have shown that LA is β-oxidized at higher rates than other fatty acids, suggesting a structural CPT1α preference for LA catalysis [77,78].

Regardless of the underlying mechanism, one question that emerges from these findings is how LA depletion affects SLD progression. Because LA can be elongated to AA, major substrate for lipid peroxidation and ferroptosis and precursor for lipid mediators such as eicosanoids [12,79], LA depletion would be expected to attenuate SLD by mitigating oxidative injury and inflammation. In contrast, dietary reestablishment of liver LA content reduced hepatic TNFα and IL-10 levels, without affecting liver steatosis or lipid peroxidation. Previous findings also support a beneficial effect of LA on SLD: higher liver LA and AA are associated with reduced inflammation, steatosis, and fibrosis [18,19], and dietary LA supplementation reduced liver steatosis [20]. Mechanistically, LA depletion may promote hepatocyte injury because LA is a major component of cardiolipins, key inner mitochondrial membrane phospholipids that regulate respiratory complex formation and stability, electron transport chain efficiency, and proton leakage [80,81]. Indeed, liver LA depletion in L-PtenKO mice occurred along with a marked reduction in cardiolipin, a finding that seems paradoxical to the increased mitochondrial respiration and oxidative capacity, but it is in line with the marked reduction in cardiolipin synthase (*Crls1*) mRNA and changes in mitochondrial morphology, including the rounded shape and disorganized cristae found in L-PtenKO livers. Interestingly, previous studies have found that steatosis and steatohepatitis in both humans and mice are associated with a marked downregulation of mitochondrial cardiolipin [82], while liver deletion of cardiolipin synthase was sufficient to cause steatosis, fibrosis, and inflammation, phenocopying many features commonly found in the steatosis-to-steatohepatitis progression [82]. Noteworthy, similarly to our findings, cardiolipin deficiency was also associated with enhanced mitochondrial respiratory capacity [82], a dissociation that may be attributed to the increase in L-PtenKO livers of proteins regulating mitochondrial dynamics and protein quality control, including total and Ser616-phosphorylated DRP1, OPA1, and LONP1 [83–85].

In contrast to the mild impact of dietary LA supplementation, pharmacological ACC and DNL inhibition robustly mitigated hepatic accumulation of DNL-related fatty acids, steatosis, inflammation, and fibrosis in L-PtenKO mice, strongly indicating that constitutive DNL activation is the main driver of steatohepatitis induced by PI3K-AKT activation. Despite reducing lipid accumulation, ACC inhibition did not affect liver mass or the low levels of lipid peroxidation, suggesting that other processes induced by PI3K–AKT signaling independently of DNL, such as hepatocyte proliferation, enhanced antioxidant capacity, and glycogen accumulation, may contribute to these phenotypes. Supporting this notion, genetic disruption of hepatic ACC1/ACC2 has been reported to increase liver mass through enhanced protein synthesis during refeeding, despite reduced hepatic lipid accumulation [86], while hepatic ACC inhibition has been shown to enhance antioxidant defenses, including NADPH and glutathione availability [87].

Although *Pten* inactivation is a recurrent signature found in SLD patients [31,32], one major limitation of our study relates to the fact that constitutive PI3K-AKT signaling promotes a temporally defined SLD progression that reproduces some (steatosis, inflammation and fibrosis), but not all features commonly found in the majority of SLD patients, including oxidative stress, hyperinsulinemia, insulin resistance and obesity [88,89]. Therefore, it is of major importance to interrogate our major findings in other mouse models of SLD.

In conclusion, steatohepatitis induced by constitutive PI3K-AKT signaling in hepatocytes occurs along with reduced lipid peroxidation and is associated with the elicitation of important lipid-related metabolic cycles, altogether promoting a robust lipidome remodeling mainly defined by enrichment of DNL-derived fatty acids in detriment of a broad depletion of essential fatty acids (LA and ALA) and cardiolipins, which chronically may be harmful helping to foment disease progression. Finally, we raised important evidence indicating that enhanced DNL and liver accumulation of DNL-related fatty acids are major culprits promoting hepatic steatosis, inflammation and fibrosis, but not the reduced lipid peroxidation and essential fatty acid depletion upon constitutive PI3K-AKT activation.

## Acknowledgments and funding

This work was supported by grants from the São Paulo Research Foundation (FAPESP #15/19530-5, 19/01763-4, 20/04159-8, 22/11234-1, and 25/04262-7) and the Brazilian National Council for Scientific and Technological Development (CNPq #303784/2022-9) to WTF, by FAPESP (#2013/07937-8 and #2024/16848-3) and CNPq (#313162/2026-3) to SM and by FAPESP (#2018/14898-2) and CNPq (#408213/2024-8) to FCM. The Tri-Carb 5110 TR Liquid Scintillation Analyzer (PerkinElmer) used in this study was acquired with support from FAPESP (#2019/12668-2). ÁSP, BFL, ÉC, TSV, JVF, ABP, NMP, EVMP, MAA-H, ACPB, LS, MM, LBG, ABC-F, BPS and TEO were recipients of fellowships from FAPESP (#19/26473-9, 22/02123-1, 21/14419-0, 26/01093-2, 20/16656-6, 23/04753-5, 24/01632-5, 24/09406-4, 23/04509-7, 24/12973-8, 24/13040-5, 23/17140-1, 24/11195-1, 24/16241-1, 25/14824-2, 21/10153-5, 23/12767-6, and 26/01093-2). LPS-J, MLEM, and SS were recipients of fellowships from the Coordenação de Aperfeiçoamento de Pessoal de Nível Superior (CAPES #88887.673950/2022-00, #88887.084152/2024-00, and #88887.251830/2026-00). EMS received fellowships from CNPq (#144006/2025-1). The authors acknowledge the use of BioRender (BioRender–biorender.com) to create the abstract graphic.

## Authors contributions

ASP and WTF conceived and designed the study, performed and supervised the experiments, analyzed and interpreted the data, and wrote and revised the manuscript. ÉC, BFL, TEO, ABC-F, TSV, CAT, NMP, EVMP, MAA-H, ABP, LPS-J, EMS, CN, MY, and BPS performed experiments, analyzed and interpreted data, and contributed to manuscript revision. FCM, MY, and SM contributed to data analysis and interpretation and critically revised the manuscript. All authors read and approved the final manuscript.

---

**Supplementary Figure 1.**
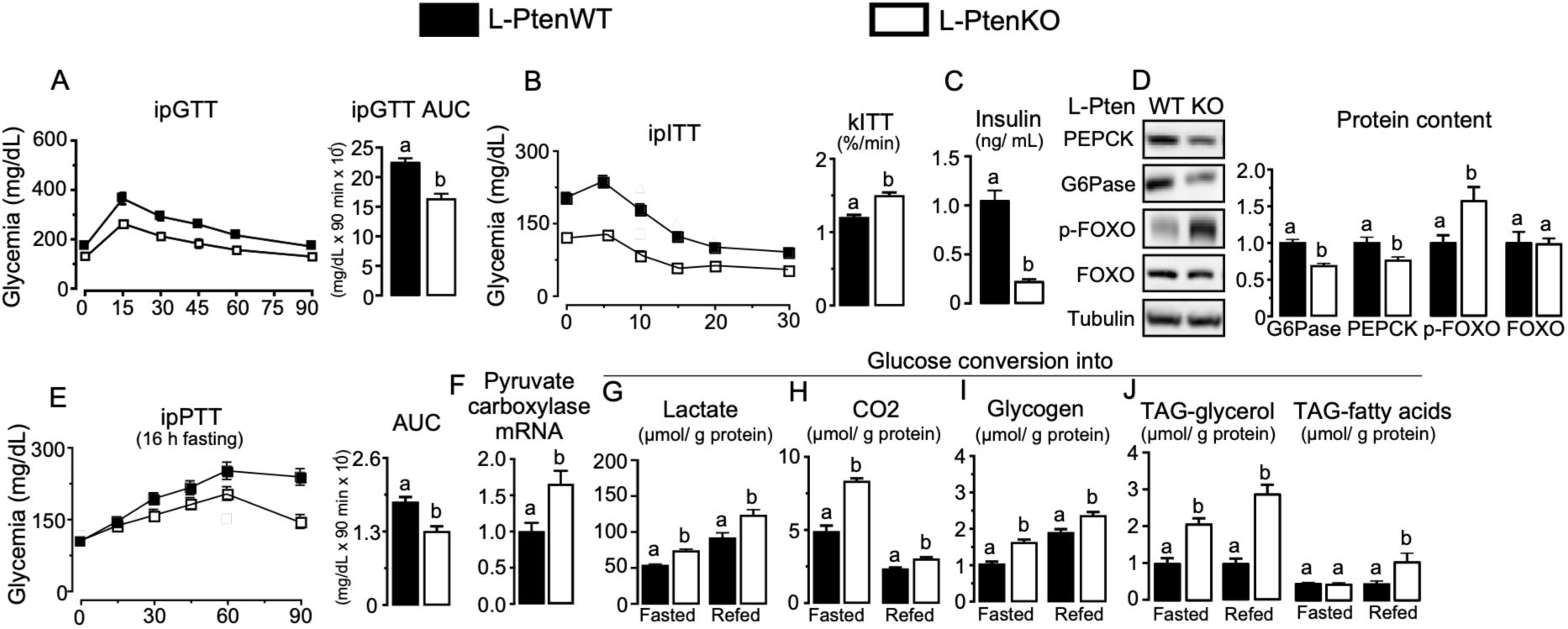
Pten deletion improves glucose tolerance and alters hepatic glucose metabolism. (A) Intraperitoneal glucose tolerance test (ipGTT) and area under the curve (AUC). (B) Intraperitoneal insulin tolerance test (ipITT), glucose disappearance rate (kITT), and serum insulin levels. (C) Intraperitoneal pyruvate tolerance test (ipPTT) and AUC after a 16-h fast. (D) Representative Western blot analysis and quantification of hepatic phosphoenolpyruvate carboxykinase (PEPCK), glucose 6-phosphatase (G6Pase), phosphorylated FOXO1 at Ser256, and total FOXO1 protein abundance. Tubulin was used as a loading control. (E) Hepatic pyruvate carboxylase mRNA expression. (F–J) Ex vivo hepatic glucose conversion into (F) lactate, (G) CO₂, (H) glycogen, (I) TAG-glycerol, and (J) TAG-fatty acids in fasted and refed mice. L-PtenWT, littermate control mice (*Pten* floxed); L-PtenKO, liver-specific *Pten* knockout mice (*Pten* floxed albumin-Cre+/−). Results are expressed as mean ± SEM. n = 5–7 mice per group. Student’s *t*-test was used to analyze the effect of *Pten* deletion (L-PtenWT vs. L-PtenKO). Means with different superscript letters differ significantly, *P* ≤ 0.05.

**Supplementary Figure 2.**
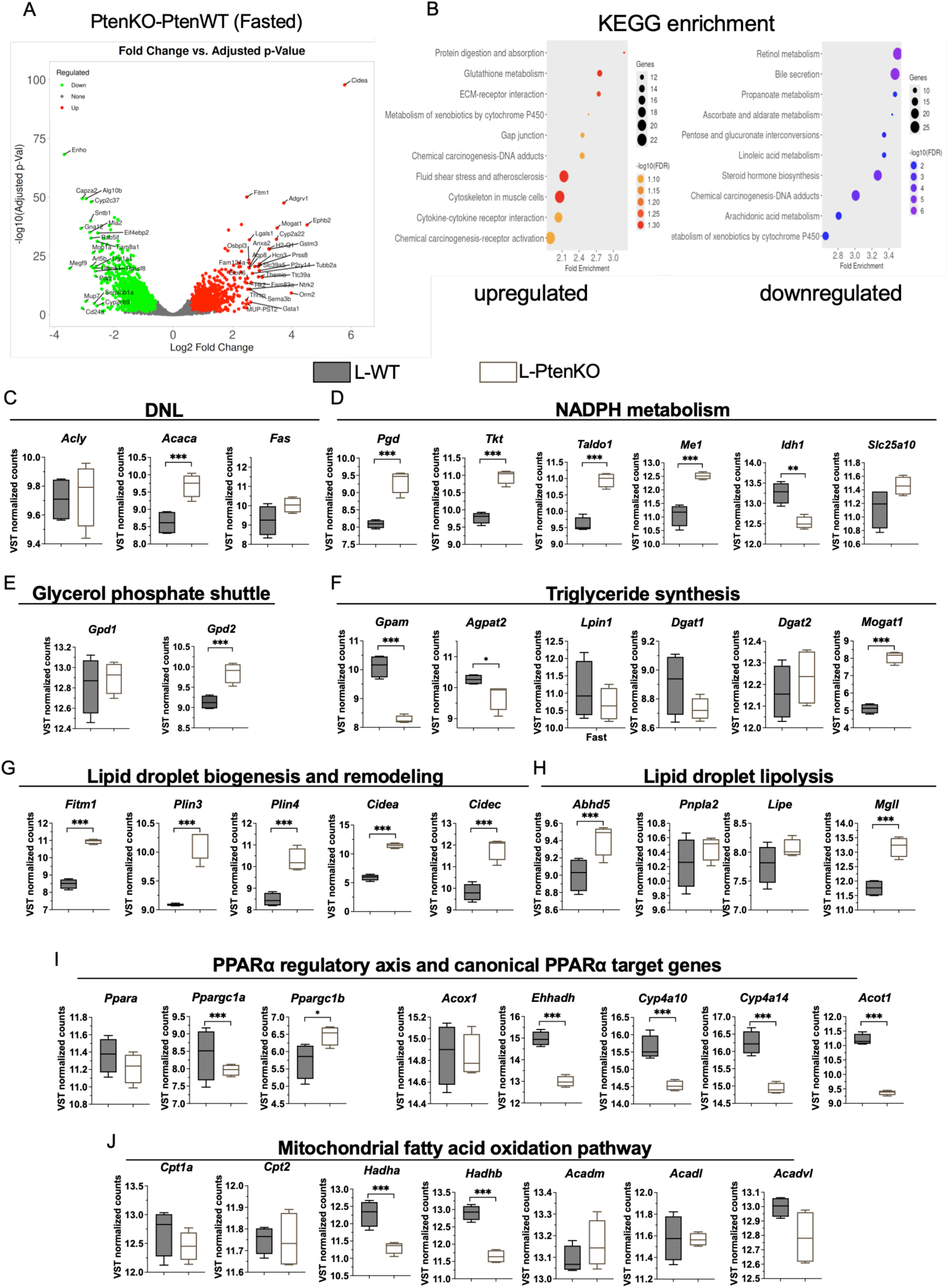
Transcriptomic remodeling following liver-specific *Pten* deletion reveals coordinated alterations in lipid and glucose metabolism. (A) Volcano plot showing differentially expressed genes between L-PtenKO and L-PtenWT livers in the fasted state. (B) KEGG pathway enrichment analysis of upregulated and downregulated genes in L-PtenKO versus L-PtenWT livers. Dot size represents the number of genes in each pathway, color indicates −log10 of the false discovery rate (FDR), and the x-axis represents fold enrichment. (C–J) VST-normalized expression of genes involved in (C) de novo lipogenesis (DNL), (D) NADPH metabolism, (E) glycerol phosphate shuttle, (F) triacylglycerol synthesis, (G) lipid droplet biogenesis and remodeling, (H) lipid droplet lipolysis, (I) the PPARα regulatory axis and canonical PPARα target genes, and (J) mitochondrial fatty acid oxidation in fasted L-PtenWT and L-PtenKO mice. L-PtenWT, littermate control mice (*Pten* floxed); L-PtenKO, liver-specific *Pten* knockout mice (*Pten* floxed albumin-Cre+/−). n = 4 mice per group. Raw RNA-seq reads were processed using Galaxy, and gene-level count data were subsequently analyzed in iDEP. Genes detected in all experimental groups were retained for analysis, and count data were variance-stabilizing transformation (VST)-normalized for visualization. Differential gene expression analysis was performed using DESeq2 with a false discovery rate (FDR) of 0.05. Genes with an absolute fold change >1.5 were considered differentially expressed. Data in (C–J) are presented as VST-normalized counts. \*\*\**P* < 0.05.

**Supplementary Figure 3.**
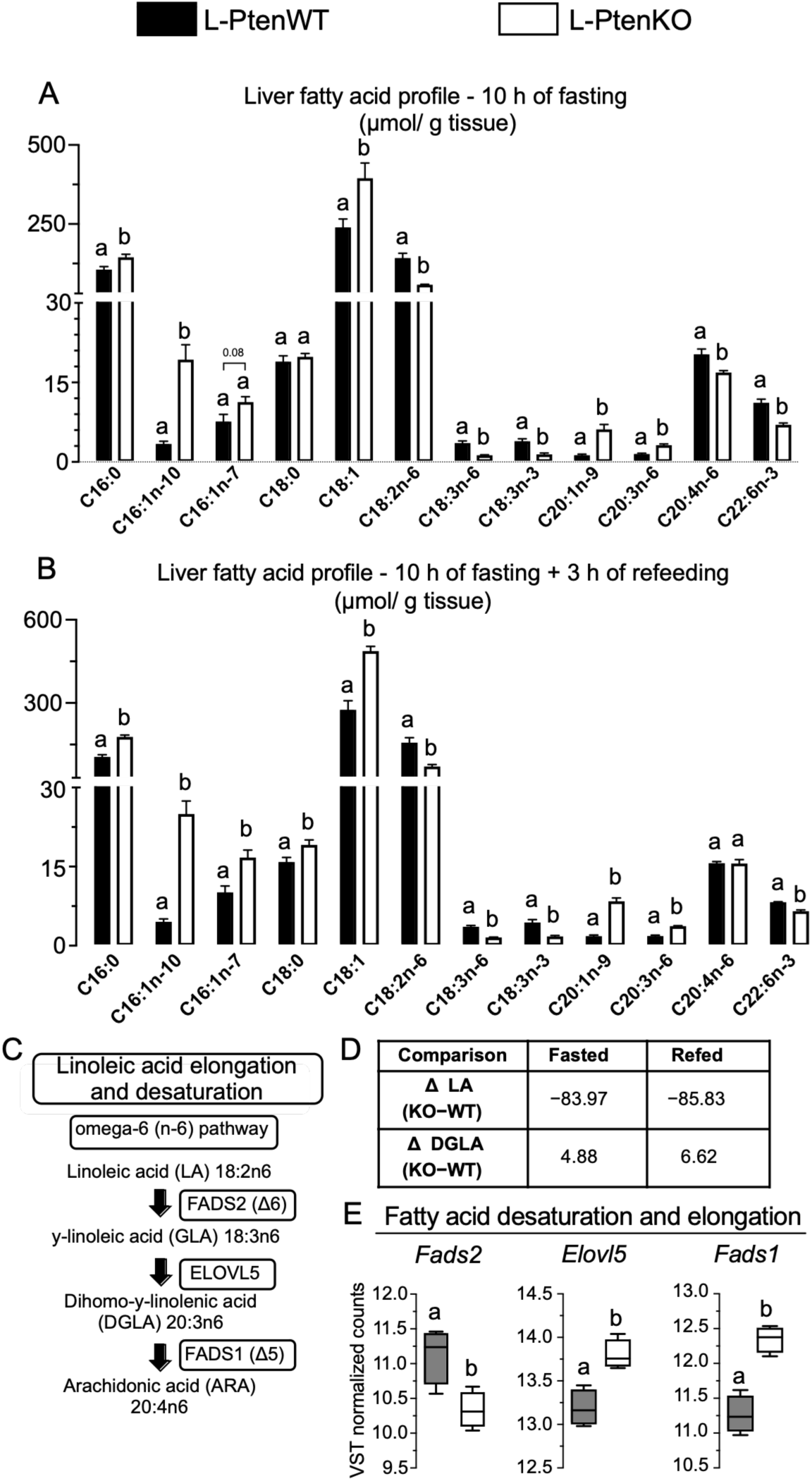
Hepatic essential fatty acid depletion following *Pten* deletion is not accompanied by increased linoleic acid elongation and desaturation. **(A–B)** Hepatic fatty acid profiles determined by GC-FAME analysis after 10 h of fasting (A) and after 10 h of fasting followed by 3 h of refeeding (B). **(C)** Schematic representation of the linoleic acid (LA) elongation and desaturation pathway, including conversion of LA (18:2n-6) to γ-linolenic acid (GLA; 18:3n-6), dihomo-γ-linolenic acid (DGLA; 20:3n-6), and arachidonic acid (AA; 20:4n-6) through FADS2, ELOVL5, and FADS1, respectively. **(D)** Differences in hepatic LA and DGLA contents between L-PtenKO and L-PtenWT mice (KO−WT) under fasted and refed conditions. **(E)** VST-normalized hepatic mRNA expression of *Fads2, Elovl5,* and *Fads1*. L-PtenWT, littermate control mice (*Pten* floxed); L-PtenKO, liver-specific *Pten* knockout mice (*Pten* floxed Albumin-Cre+/−). Results are expressed as mean ± SEM. *n* = 4–5 mice per group for GC-FAME analysis and *n* = 4 mice per group for RNA-seq analysis. Statistical significance in (A–B) was determined using Student’s *t*-test. Differential gene expression analysis in (E) was performed using DESeq2 with a false discovery rate (FDR) of 0.05. Means with different superscript letters are significantly different, *P* ≤ 0.05.

**Supplementary Figure 4.**
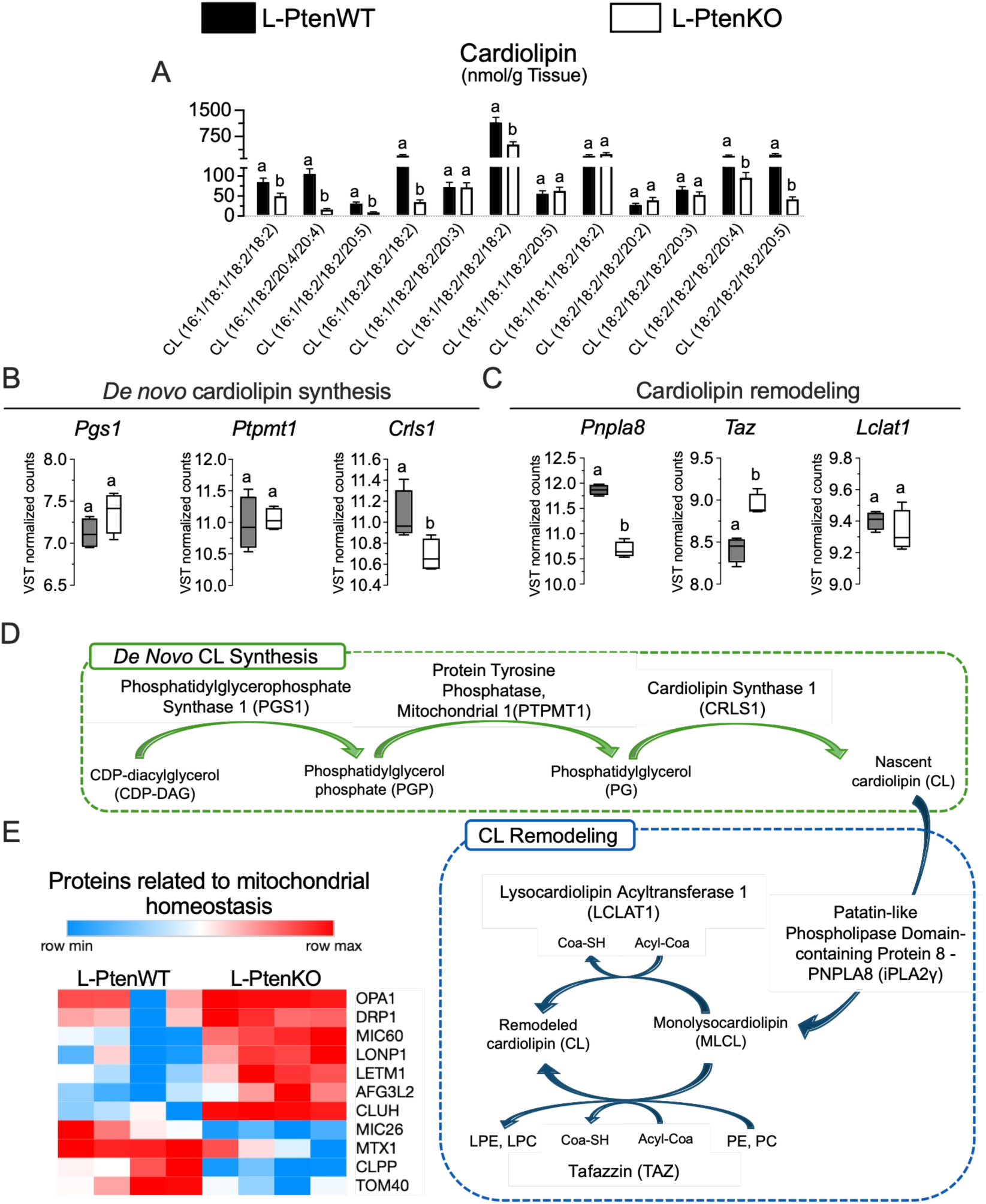
Altered cardiolipin composition following *Pten* deletion is associated with changes in cardiolipin synthesis and remodeling. **(A)** Hepatic levels of individual cardiolipin (CL) species. **(B)** VST-normalized hepatic mRNA expression of genes involved in *de novo* CL synthesis, including *Ptgs1, Ptpmt1,* and *Crls1*. **(C)** VST-normalized hepatic mRNA expression of genes involved in CL remodeling, including *Pnpla8, Taz,* and *Lclat1*. **(D)** Schematic representation of de novo CL synthesis and CL remodeling pathways. **(E)** Heatmap showing the relative abundance of proteins related to mitochondrial homeostasis in L-PtenWT and L-PtenKO livers. Protein abundance is represented by row-scaled Z-scores. L-PtenWT, littermate control mice (*Pten* floxed); L-PtenKO, liver-specific *Pten* knockout mice (*Pten* floxed Albumin-Cre+/−). *n* = 7–9 mice per group for lipidomic analysis and *n* = 4 mice per group for proteomic and RNA-seq analyses. Statistical significance in (A) was determined using Student’s *t*-test. Differential gene expression analysis in (B–C) was performed using DESeq2 with a false discovery rate (FDR) of 0.05. Means with different superscript letters are significantly different, *P* ≤ 0.05.

**Supplementary Table 1.** Low fat (LF) and high-fat diets containing either 41 (HF) or 77.3 (HFLA) g /kg compositions.

| **Ingredient (g)** | **LF** | **HF** | **HFLA** |
| --- | --- | --- | --- |
| Choline | 1.9 | 2.5 | 2.5 |
| L-Cystine | 2.85 | 1.8 | 1.8 |
| Vitamin Mix | 10.075 | 10 | 10 |
| Mineral Mix | 35.25 | 35 | 35 |
| Cellulose | 55 | 50 | 50 |
| Sucrose | 89.55 | 100 | 100 |
| Maltodextrin 10 | 162.675 | 100 | 100 |
| Casein | 190.95 | 220 | 220 |
| Corn Starch | 416.05 | 273.55 | 273.55 |
| Soybean oil | 23.85 | 40 | 40 |
| Pure Linoleic Acid | 0 | 0 | 41.4 |
| Porc Lard | 19.1 | 167 | 125.6 |
| Vitamin E | 0.15 | 0.15 | 0.15 |
| Water (L) | 0.25 | 0 | 0 |
| total | 1007.65 | 1000 | 1000 |
| **Energy content** |  | **Kcal** |  |
| Protein | 775.2 | 887.2 | 887.2 |
| Carbohydrate | 2673.1 | 1894.2 | 1894.2 |
| Fat | 386.55 | 1863 | 1863 |
| Total Kcal | 3834.85 | 4644.4 | 4644.4 |
| **Macronutrient distribution** |  | **% Kcal** |  |
| Protein | 20.2 | 19.1 | 19.1 |
| Carbohydrate | 69.7 | 40.8 | 40.8 |
| Fat | 10.1 | 40.1 | 40.1 |
| Total Kcal | 100.0 | 100.0 | 100.0 |
| **Fatty acid composition** |  | **(g)** |  |
| Linoleic Acid C18:2) | 14.4 | 41.0 | 77.3 |
| Palmitic Acid (C16:0) | 7.0 | 62.92 | 46.7 |
| Oleic Acid (C18:1) | 13.8 | 116.38 | 87.1 |

**Supplementary Table 2.**
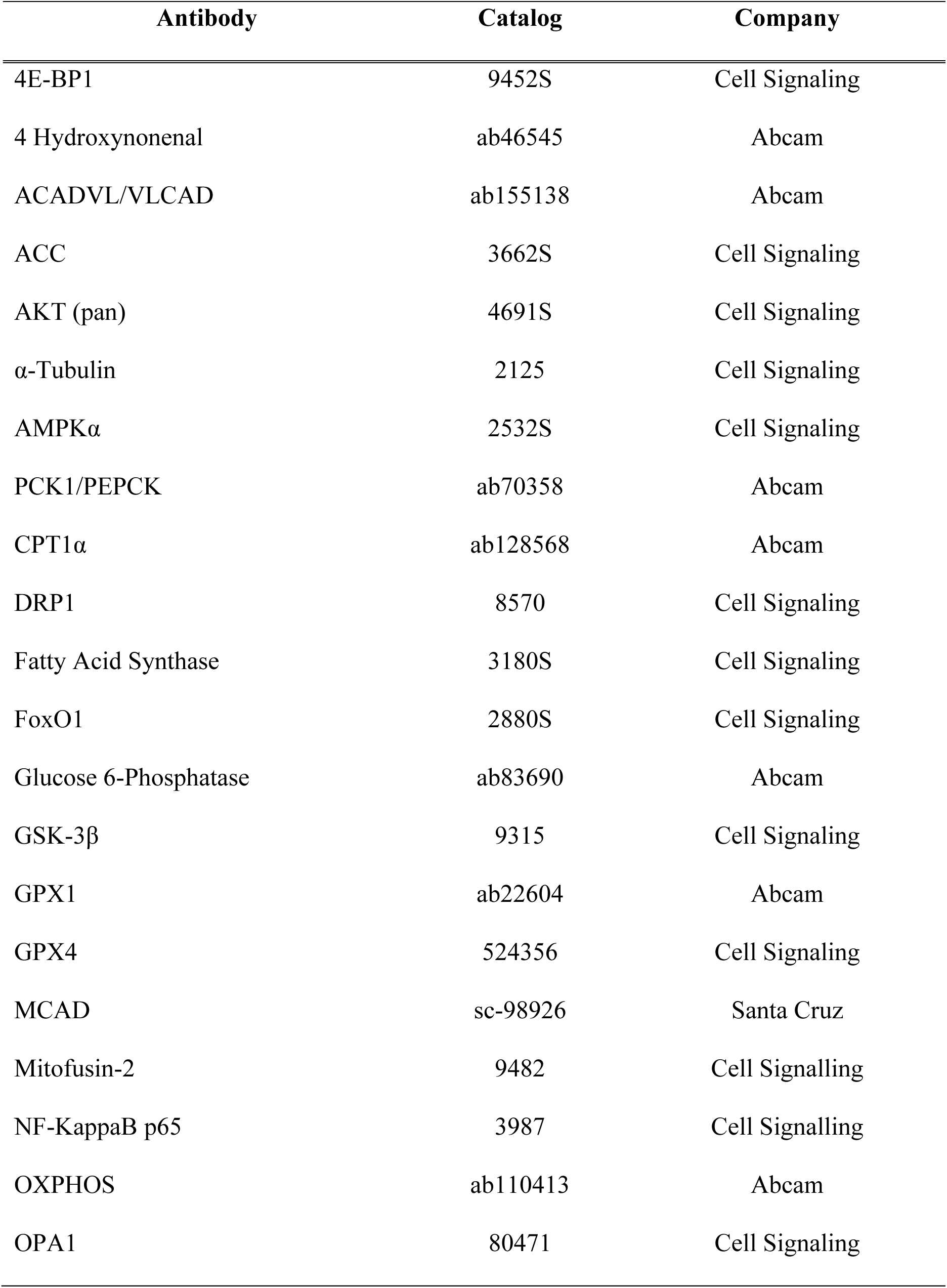
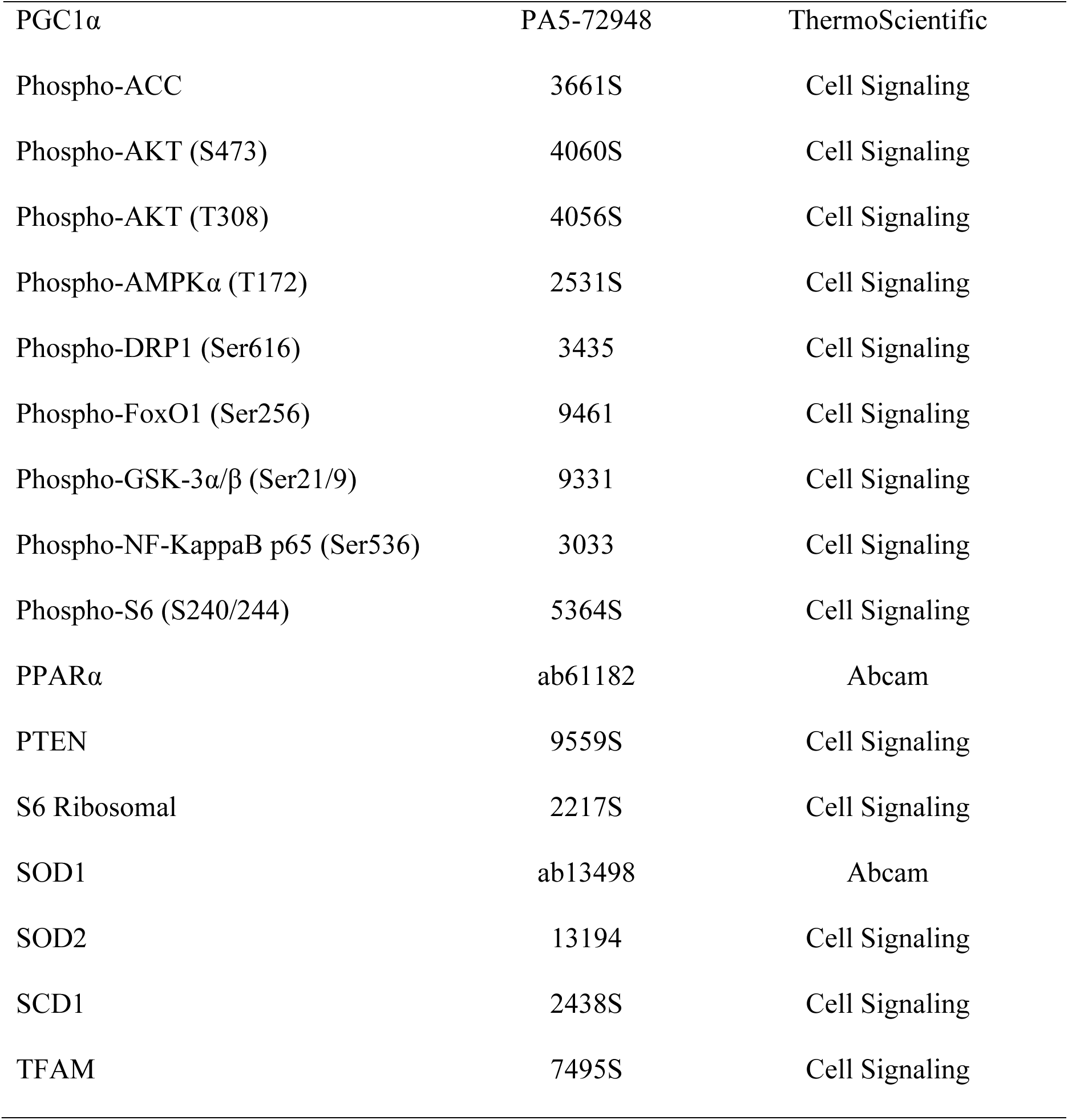
List of antibodies used.

**Supplementary Table 3.**
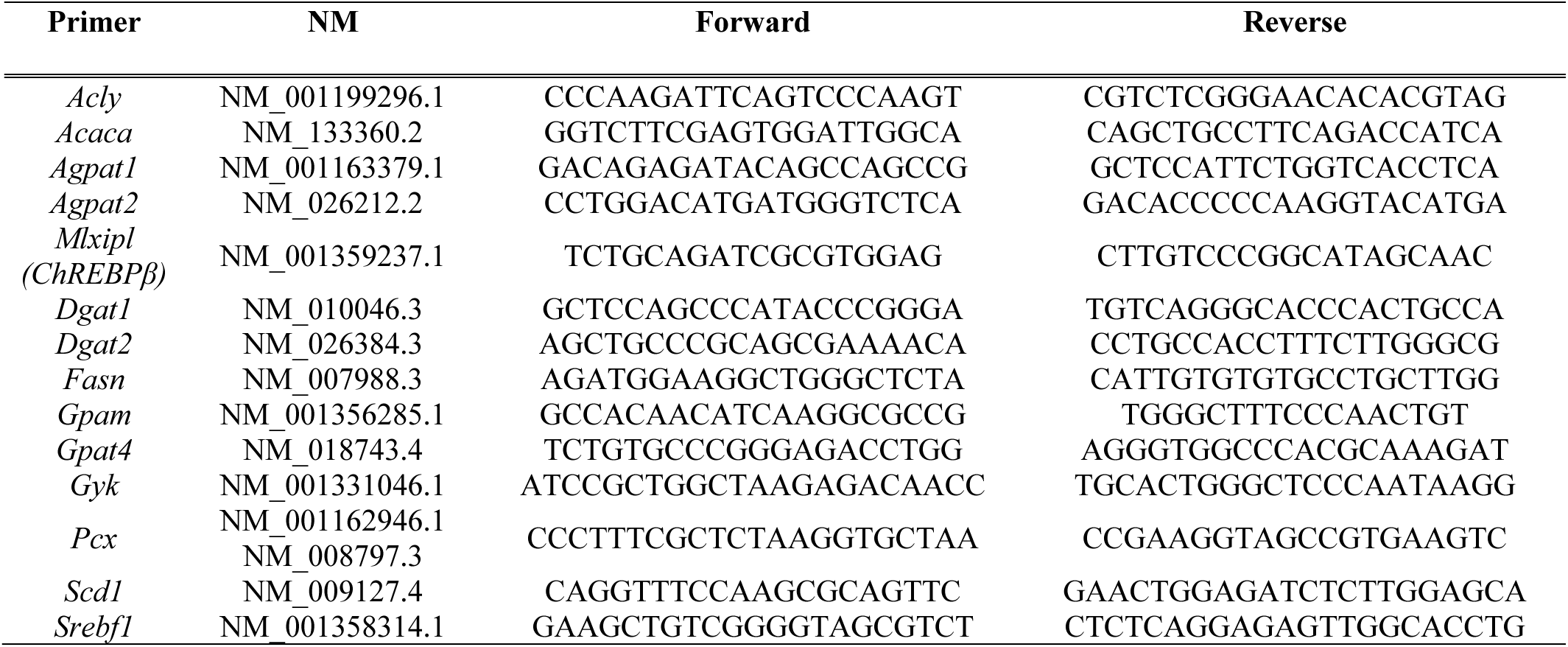
List of primers used for RT-PCR.

**Supplementary Table 4.**
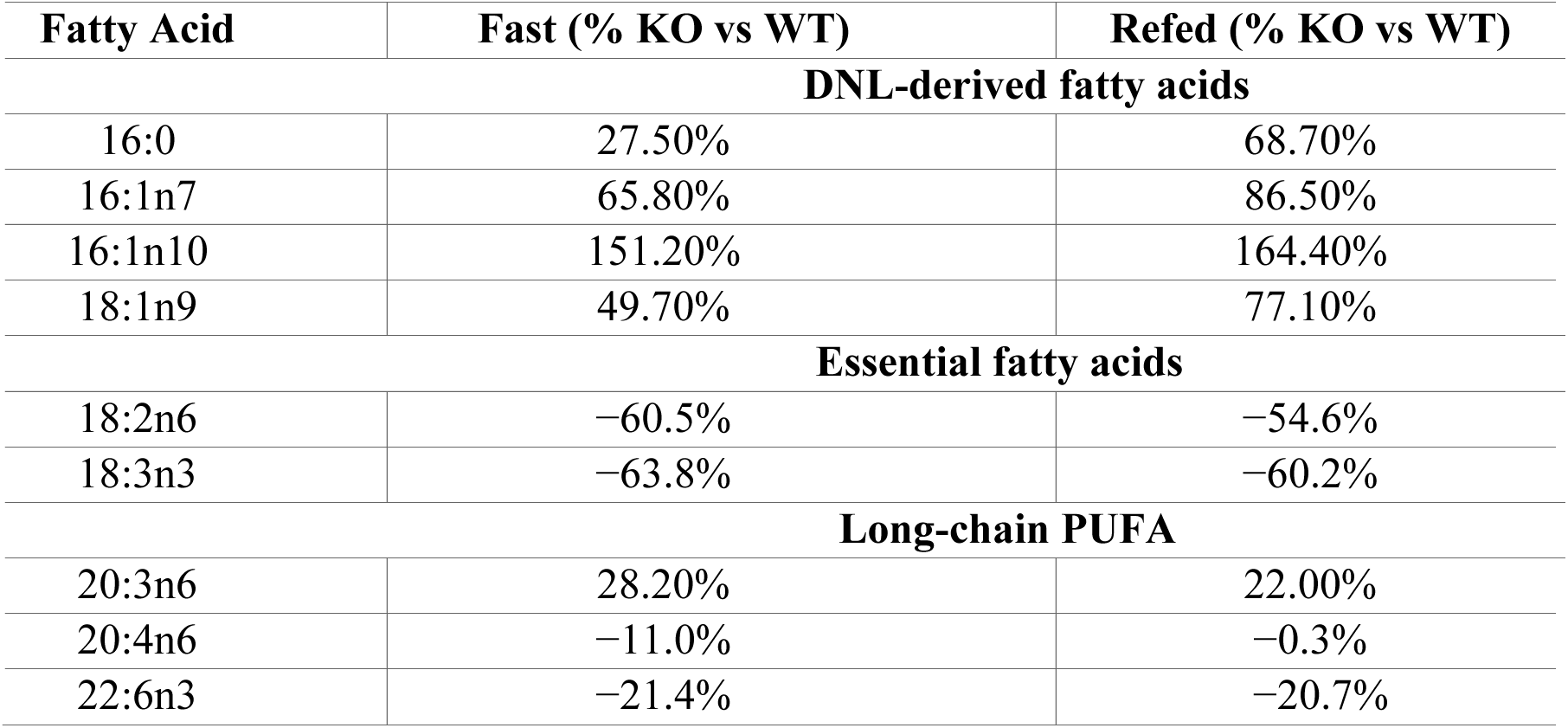
Differential effects of hepatocyte Pten deletion on DNL-derived and essential fatty acid pools in liver.

**Supplementary Table 5.** Fasting-refeeding regulation of DNL-derived and essential hepatic fatty acid pools in L-PtenKO mice.

| **Fatty Acid** | **WT Refed/Fast (%)** | **KO Refed/Fast (%)** |
| --- | --- | --- |
| **DNL-derived fatty acids** |  |  |
| 16:0 | 68.20% | 85.80% |
| 16:1n7 | 44.40% | 87.50% |
| 16:1n10 | 33.60% | 119.10% |
| 18:1n9 | 15.10% | 36.30% |
| **Essential fatty acids** |  |  |
| 18:2n6 | 10.40% | 26.20% |
| 18:3n3 | 13.00% | 24.70% |
| **Long-chain PUFA** |  |  |
| 20:3n6 | 55.30% | 50.10% |
| 20:4n6 | 10.20% | 21.40% |
| 22:6n3 | -26.2% | -19.2% |

